# Low concentrations of ethylene bisdithiocarbamate pesticides maneb and mancozeb impair manganese and zinc homeostasis to induce oxidative stress and caspase-dependent apoptosis in human hepatocytes

**DOI:** 10.1101/2023.07.11.548490

**Authors:** Kilian Petitjean, Yann Verres, Sébastien Bristeau, Catherine Ribault, Caroline Aninat, Christophe Olivier, Patricia Leroyer, Martine Ropert, Olivier Loreal, Olivier Herault, Laurence Amalric, Nicole Baran, Bernard Fromenty, Anne Corlu, Pascal Loyer

**Author notes:** **Contact information**: LOYER Pascal, NuMeCan Inserm UMR 1317, CHU Pontchaillou, 35033 Rennes, France.; CORLU Anne, NuMeCan Inserm UMR 1317, CHU Pontchaillou, 35033 Rennes, France. **Financial support**: This work was funded by the Cancéropole Grand-Ouest (CGO projet structurant PeNiCa), Institut National de la Santé et de la Recherche Médicale (Inserm, France) and the “Organisme Française pour la Biodiversité (OFB), programme ECOPHYTO II”, project PESTIFAT AFB/2019/87 (PR-EST, 2019-2022). Kilian Petitjean received fellowships from the CGO (PeNiCa, 2018) and the OFB (PESTIFAT, 2019-2022).

## Abstract

The worldwide and intensive use of phytosanitary compounds results in environmental and food contamination by chemical residues. Human exposure to multiple pesticide residues is a major health issue. Considering that the liver is not only the main organ for metabolizing pesticides but also a major target of toxicities induced by xenobiotics, we studied the effects of a mixture of 7 pesticides (chlorpyrifos-ethyl, dimethoate, diazinon, iprodione, imazalil, maneb, mancozeb) often detected in food samples. Effects of the mixture was investigated using metabolically competent HepaRG cells and human hepatocytes in primary culture. We report the strong cytotoxicity of the pesticide mixture towards hepatocytes-like HepaRG cells and human hepatocytes upon acute and chronic exposures at low concentrations extrapolated from the Acceptable Daily Intake (ADI) of each compound. Unexpectedly, we demonstrated that the manganese (Mn)-containing dithiocarbamates (DTCs) maneb and mancozeb were solely responsible for the cytotoxicity induced by the mixture. The mechanism of cell death involved the induction of oxidative stress, which led to cell death by intrinsic apoptosis involving caspases 3 and 9. Importantly, this cytotoxic effect was found only in cells metabolizing these pesticides. Herein, we unveil a novel mechanism of toxicity of the Mn-containing DTCs maneb and mancozeb through their metabolization in hepatocytes generating the main metabolite ethylene thiourea (ETU) and the release of Mn leading to intracellular Mn overload and depletion in zinc (Zn). Alteration of the Mn and Zn homeostasis provokes the oxidative stress and the induction of apoptosis, which can be prevented by Zn supplementation. Our data demonstrate the hepatotoxicity of Mn-containing fungicides at very low doses and unveil their adverse effect in disrupting Mn and Zn homeostasis and triggering oxidative stress in human hepatocytes.

## 1. Introduction

Pesticides remain extensively used in agricultural practices despite the well-documented risks for the human health (Alvanja, 2009; Deziel et al., 2017). Occupational exposures to pesticides have been clearly correlated to higher risk of a wide range of pathologies including developmental abnormalities (Rauh et al., 2012), Parkinson disease (Brouwer et al., 2017; Pouchieu et al., 2018), hematological disorders (Merhi et al., 2007; Jin et al., 2014), solid tumors (Sabarwal et al., 2018; Lerro et al., 2021) and leukemia (Ferri et al., 2017; Foucault et al., 2021). Higher occurrences of pesticide-dependent diseases are not restricted to occupational exposure but are also documented in populations living in agricultural areas of intensive spreading via the drifting of pesticides in the air and house dust contamination (Deziel et al., 2017).

The intensive use of pesticides contributes to the so-called diffuse pollution (Mghirbi et al., 2018) with accumulation of chemicals in the upper layers of soil as a result of atmospheric deposition (Mali et al., 2023) and land application (Ritter et al., 2002) of these compounds. The soil contamination and the direct spreading of pesticides on crops result in the presence of chemical residues in agricultural products (Czaja et al., 2020). Regulatory authorities such as the Food and Drug Administration (US FDA), the European Food Safety Authority (EFSA) and the Organization for Economic Co-operation and Development (OECD) defined residues as the mixture of active substances, metabolites and degradation products and implemented the Maximum Residue Levels (MRLs) corresponding to the highest levels of pesticide residues legally tolerated in agricultural commodities when pesticides are applied correctly.

These agencies also established dietary risk assessments of general population to pesticide exposures by monitoring the presence of residues in food samples (OECD 2009; EFSA 2017; US FDA 2019) and considering the toxicity of each contaminant as well as the overall amounts used for each pesticide. In the past decade, the presence of at least one chemical contaminant in nearly 50% of food products and multiple residues in 25% of the samples were reported (OECD 2009; EFSA 2017; US FDA 2019). In most cases, the amounts of residues in human foods were compliant with MRLs established standards, however, 10% of these commodities showed contamination levels that exceeded those maximum accepted levels. Considering the large diversity of active substances used in agricultural practice, it is practically impossible to monitor all active substances and their metabolites. For instance, the dithiocarbamates (DTCs) fungicides maneb and mancozeb used worldwide are poorly water soluble and bind tightly to soil. Because of their unstable polymeric structures in aqueous solution, maneb and mancozeb are difficult to identify by analytical procedures. Their hydrolysis generates fragmented complexes further metabolized by plants and animals in ethylene thiourea (ETU), their main metabolite. In absence of analytical protocols to analyze each DTC independently and all their residues, total DTC concentrations can be measured in water (Bristeau and Ghestem, 2018; Bristeau and Amalric, 2021). Nevertheless, the assessment of contamination levels in commodities remains poorly characterized and exposure to DTCs in humans relies only on the detection of ETU in urine (Stadler et al., 2022).

At the European level, the approval of pesticides is based on individual toxicology studies of the active substance. The EFSA established for each pesticide an Acceptable Daily Intake (ADI), which is a theoretical safety threshold for human health calculated from the No observable Adverse Effect Level (NoAEL) determined in animal models applying a corrective factor to account for intra-species and intra-individual variations (Dorne, 2010). Nevertheless, ADI does not consider potential cumulative effects of pesticide mixtures while regulatory agencies recognized that consumers are exposed to multiple residues present in food for long periods of times, which could lead to cumulative and delayed effects on human health. In this context, it is necessary to test low doses of mixtures of pesticides belonging to different families with similar or different mechanisms of action to better assess the possible existence of a risk to human health. This combined exposure could lead to insidious synergistic effects significantly different from those observed if summing the effect of each pesticide evaluated individually.

In order to extend knowledge on cellular alterations induced by pesticide mixtures, the consortium project entitled “Pesticides and Tumor Niches” (Cancéropole Grand-Ouest, France) studied the effects of a mixture of 7 pesticides from different classes of chemicals (chlorpyrifos-ethyl, dimethoate, diazinon, iprodione, imazalil, maneb, mancozeb) often detected in food samples (EFSA, 2017). These compounds were used at concentrations extrapolated from the ADI dose for each pesticide in various *in vitro* cell models. In bone marrow mesenchymal stromal cells (BM-MSCs), it was shown that the pesticide mixture triggered the senescence of BM-MSCs, their differentiation in adipocytes (Hochane et al., 2017) and the deterioration of their immunosuppressive properties (Leveque et al., 2019). These pesticides also altered the BM-MSC capacity to support primitive hematopoiesis by monitoring *in vitro* expansion of human normal BM CD34^+^ progenitors co-cultured with pesticides-modified BM-MSCs (Foucault et al., 2021). In this last study, the manganese (Mn)-containing DTCs, maneb and mancozeb significantly contributed to the decrease in the CD34+ cell expansion (Foucault et al., 2021). Interestingly, similar alterations have been described in BM-MSCs of patients suffering from myelodysplastic syndromes supporting the hypothesis that exposure to pesticides could favor the appearance of pre-leukemic states in the elderly (Foucault et al., 2021).

Most pesticides are metabolized in the liver by phase I and II enzymes to produce hydrophilic products more easily eliminated in urine and bile (Jandacek and Tso, 2007; Klimowska et al., 2020; Kernalléguen et al., 2022). The liver is also a major target for toxicities triggered by xenobiotics themselves and/or their downstream reactive metabolites. In the present study, we evaluated the effects of the pesticide mixture and the 7 pesticides tested individually on metabolically competent hepatocytes *in vitro* at concentrations that previously showed effects of BM-MSCs. We report that the pesticide mixture induced a strong cytotoxicity in cultures of hepatocyte-like HepaRG cells and human hepatocytes following acute and chronic exposures at low concentrations extrapolated from the ADI. In addition, we demonstrated that the metal-containing DTCs maneb and mancozeb were solely responsible for the cytotoxicity induced by the mixture but only in cells metabolizing these fungicides through the intracellular accumulation of Mn and reduction in zinc (Zn) content, which triggered oxidative stress and apoptosis. Together, our data unveil a novel mechanism of hepatotoxicity of Mn-containing fungicides and emphasize the crucial role of the Mn and Zn homeostasis to regulate ROS production and cell fate in human hepatocytes.

## 2. Materials and methods

### 2.1. Pesticides, mixture, doses and treatments

Chlorpyrifos, dimethoate, diazinon, imazalil, iprodione, maneb, mancozeb, zineb Pestanal^TM^ analytical standards, ethylenethiourea (45531), ethyleneurea (31534) and ethylenediamine (41008) were purchased from Sigma Aldrich (St. Louis, MO, USA). CAS numbers and chemical structures are provided in **Supporting Information 1**.

The concentrations of pesticides used to treat the cells were extrapolated from two values (Hochane et al., 2017; Leveque et al., 2019): 1) the international ADI representing the threshold of safety in humans for lifetime exposure. Values of ADI (mg/kg.bw/day) for each pesticide was converted to g/L and then in mol/L for *in vitro* models, 2) the high nutritional daily intake (hNDI) or Theorical Maximum Daily Intake (TMDI) corresponding to the lowest concentration of pesticides used in our study. Values of hNDI further referred as percentages of ADI (%ADI) correspond to the real exposure of the French population of all ages for each pesticide and calculated with the MRLs in food commodities established by EFSA. For the values of hNDI (%ADI), ADI, and 3ADI, a conversion has been made in order to extrapolate the aforementioned doses (estimated in mg/kg body weight/day) to an *in vitro* model. We considered a total absorption of this ingested amount and then its dilution in 5 L of blood in a subject of 60 kg, in order to obtain the blood concentration (mg/L and μmol/L) to which the various organs could be theoretically exposed.

Dimethyl-sulfoxide (DMSO) was the solvent used to prepare stock solution at 100mM except for the diazinon at 10mM. These stock solutions were diluted in culture medium to obtain the final concentrations of exposure. Control condition corresponded to cultures treated with the same concentration of DMSO without pesticides. For differentiated HepaRG cells and human hepatocytes, DMSO in the culture medium was adjusted at 2% (vol/vol), which corresponds to the culture medium used routinely for keeping these cells differentiated. DMSO concentration was 0.1% for progenitor HepaRG cells. For acute exposures cells were treated for 16 to 48 hours depending on the assays performed after exposure. For chronic exposures, cells were treated at each culture medium renewal, 3 times a week.

### 2.2. Cell cultures

The human hepatocellular HepaRG progenitor cells actively proliferate at low density and are capable to differentiate into biliary- and hepatocyte-like cells (Gripon et al., 2002) in appropriate culture conditions. Progenitor HepaRG cells were cultured as previously described (Cerec et al., 2007) in William’s E Medium supplemented with Fetal Bovine Serum (FBS) 10% (Eurobio batch S52446-2170, reference CVFSVF00-01, HyClone Fœtal II batch AAF204244 reference SH30066.03, ratio 1-1), 1% glutamine (200mM), 100U/ml penicillin, 100µg/ml streptomycin, 5µg/ml insulin, and 50µM hydrocortisone hemisuccinate (Upjohn pharmacia). HepaRG cells were seeded at a density of 2.6×10^4^ cells/cm^2^. After 2 weeks, cells were cultured in the same medium supplemented with 2% DMSO to obtain differentiated HepaRG cells including hepatocyte- and cholangiocyte-like cells (Cerec et al., 2007). In some experiments, hepatocyte- and cholangiocytes-like HepaRG cells were selectively detached using mild trypsinization with Trypsin 0.05% diluted in PBS (vol/vol) as previously described (Cerec et al., 2007; Laurent et al., 2013) and plated separately at high density (2.5×10^5^ cells/cm^2^) to obtain cultures enriched in hepatocytes or cholangiocytes. Both hepatocyte- and cholangiocyte-like cells were cultured in medium supplemented with 2% DMSO.

Human liver biopsies and primary hepatocytes were obtained from the processing of biological samples through the Centre de Ressources Biologiques (CRB) Santé of Rennes (Institutional review board approval BB-0033-00,056) under French legal guidelines and fulfilled the requirements of the institutional ethics committee. Primary human hepatocytes (PHH) were seeded at a density of 1.5×10^5^ cells/cm^2^ and cultured in same medium than differentiated HepaRG hepatocytes. Culture procedures of human macrophages and BM-MSC, as well as rat liver epithelia cells SDVI, Huh7 and HepG2 hepatoma cells are described in **Supporting Information 1**.

### 2.3. Cell viability, apoptosis assessment and genotoxicity

Cell viability was assessed by measuring the relative intracellular ATP content using CellTiter-Glo Luminescent Cell viability assay (Promega). After treatment with pesticides, cells were incubated with the CellTiter-Glo reagent for 10 minutes (min). Lysates were transferred to an opaque multi-well plate and luminescent signals were quantified at 540nm using the Polarstar Omega microplate reader (BMG Labtech). Cell viabilities in treated cells were expressed as the percentage of the luminescent values obtained in untreated cells, which was arbitrary set as 100%. Loss of plasma membrane integrity and cytotoxicity during treatments were assessed by measuring the lactate dehydrogenase (LDH) release in the culture medium using Pierce LDH Cytotoxicity Assay Kit (Thermo Scientific) according to manufacturer’s instructions.

Caspases activities were quantified after 24 or 48 hours of treatment for HepaRG cells and PHH, respectively. Cells were lysed with the “caspase-activity” buffer containing PIPES (20mM: pH 7.2), NaCl (100mM), EDTA (1mM), CHAPS (0.1%), sucrose (10%). Lysates were homogenized using sonication for 10 seconds twice. Then, 50µg proteins were added to the caspase-activity buffer supplemented with 10mM DTT and 80μM of substrates Ac-DEVD-AMC, Ac-IEPD-AMC and Ac-LEHD-AMD (Enzo Life Science) to measure caspase 3, 8 and 9 activities, respectively. The luminescence was measured using the Polarstar Omega microplate reader (BMG Labtech) with excitation wavelengths at 380 and emission at 440nm each 10 min during a 2h-time course.

The γ-H2AX test was performed to assess genotoxicity. Briefly, cells were rinsed with phosphate-buffered saline (PBS), fixed with 4% formaldehyde in PBS for 15 min at room temperature before permeabilization using 0.1% saponin in PBS. Nonspecific binding was blocked for 15 min by incubating the cells with 2 % FBS in PBS. Then, the cells were incubated with rabbit polyclonal anti-phospho-H2AX antibody (Abcam, ab88671) diluted 1/500 in 2 % FBS-PBS solution for 1h at room temperature. Following washes with PBS, Dylight 549-labeled secondary antibody in 2 % FBS-PBS solution containing Hoechst 33342 dye (1μg/mL) was incubated for 45 min. After washing, the γ-H2AX positive nuclei were observed and ten pictures per well were analyzed corresponding to at least 1000 cells scored.

### 2.4. Detection of reactive oxygen species (ROS)

Cell-permeant CellROX Orange (Thermo Fisher Scientific, C10493) and MitoSOX (Thermo Fisher Scientific, M36008) molecular probes were used to assess ROS generation. Cells were washed with prewarmed (37°C) Hanks’ Balanced Salt Solution (HBSS) and incubated with CellROX Orange (500nM) or MitoSOX (2µM) diluted in HBSS for 30 min at 37°C, 5% CO_2_ protected from light. Then, cells were rinsed with warm HBSS. N-acetylcysteine (NAC) at 20mM was used to neutralize ROS by treating cells for 2 hours prior to and during the treatment with pesticides. Treatments with excess of zinc chloride (ZnCl_2_) at 70µM were also performed during pesticide and manganese chloride (MnCl_2_) exposures. For MitoSOX probe, fluorescence intensity was measured using the Polarstar Omega microplate reader with excitation/emission wavelengths and 520/590nm. Fluorescence intensity values were normalized to amounts of total proteins. For CellROX Orange probe, the fluorescence intensity of at least 5,000 cells was analyzed with a Becton Dickinson LSRFortessa™ X-20 (cytometry core facility of the Biology and Health Federative research structure Biosit, Rennes, France). Data were expressed as the percentage of fluorescence signal measured in untreated control cells, which was arbitrarily set as 100%.

### 2.5. Measurement of mitochondrial respiration and oxidative phosphorylation (OXPHOS) complex enzyme activities

Parameters of mitochondrial respiration were analyzed using XF Cell Mito Stress Test Kits and Agilent Seahorse XFe24 analyzer according to manufacturer’s instructions. Briefly, after 16 hours of exposure to pesticides, HepaRG cells were incubated with Seahorse XF DMEM Medium (pH 7.4) at 37°C for 1 hour. Then, Seahorse XF DMEM Medium (pH 7.4) was replaced by the same medium containing L-glutamine (2mM), glucose (10mM) and pyruvate (1mM). HepaRG cells were transferred in Agilent Seahorse XFe24 analyzer for assessment of mitochondrial respiration. Oxygen consumption rates corresponding to basal respiration, maximal respiration, ATP production and proton leak were normalized by fluorescence intensity of Hoechst 33342 dye measured with the Polarstar Omega microplate reader (BMG Labtech) with excitation/emission wavelengths at 355nm/460nm.

In order to measure OXPHOS complex enzyme activities mitochondria were first isolated from HepaRG cells treated for 16h with maneb at 6ADI, MnCl_2_ at 14µM or control solvent vehicle. Briefly, cells were harvested and washed with cold PBS, then cells were centrifuged and pellets were frozen in liquid nitrogen. Cells were resuspended within isolation buffer pH 7.2 containing mannitol (220mM), sucrose (75mM), Trizma (10mM) and EDTA (1mM). After 3 min frozen in liquid nitrogen, cells were thawed at 37°C, centrifuged 1min at 20,000g and finally resuspended in the isolation buffer. For complexes I and V and citrate synthase cells were sonicated (6 times for 5 s).

The activities of the 5 mitochondrial OXPHOS complex enzymes were then measured at 37°C on a UVmc^2^ spectrophotometer (SAFAS, Monaco) as previously described (Chrétien et al., 2004; Medja et al., 2009) with specific buffer and reagents for each complex (**Supporting Information 1**). Absorbance changes due to the respective substrate conversions were analyzed at, 600nm for complex I and II, 550nm for complex III and IV, 340nm for complex V and 412nm for citrate synthase. Enzymes activities of all complexes were expressed as nmol substrate/min/10^6^cells using the Beer Lambert’s law and were normalized to the citrate synthase activity.

### 2.6. Studies of mancozeb metabolism by High pressure liquid chromatography (HPLC) and liquid chromatography coupled to mass spectrometry (LC-MS/MS)

The HPLC analysis was performed with the Agilent HPLC 1260 Infinity II with the column Accucore PFP (Thermo Scientific ref: 17426-153030) coupled with a UV detector. Cells were washed with PBS and incubated with phenol red-free William’s E Medium containing 100µM, maneb or mancozeb. After incubations of cells for 3, 8 or 24 hours, the medium was collected and centrifuged 15 min at 20,000g at 4°C. 50µL of supernatant was injected in HPLC system. A gradient of acetonitrile and 0.1% acetic acid was used during the run at a flow rate of 0.500 mL/min. The run starts at 100% of acetic acid and decrease to 10% in 20 min. After 2 min at 10% acetic acid and 90% acetonitrile, the gradient returns to 100% acetic acid in 5 min. The wavelengths 247 and 285 nm were used to detect the metabolites of mancozeb, and the mancozeb, respectively. The peaks of mancozeb and ETU were identified by comparing the chromatograms of culture media containing or not these compounds and using analytical standards.

The analysis was performed by LC-MS/MS detection with direct injection using Xevo-TQD ® triple quadrupole UPLC/MSMS (Waters) in MRM (Multiple Reaction Monitoring) mode. The mass spectrometer was operated in the positive and negative electron ionization modes for identification of the unknown compound resulting from mancozeb and MnCl_2_ exposures while the mode of ionization positive electrospray was selected for quantification of mancozeb metabolites. The injection of analytical standards allowed to define the analytical parameters of the 8 metabolites and active substance (**Supporting Information 2**) and to estimate their quantification limits. To investigate the linearity of the method, 9 concentrations (0.01 to 10µM in a mixture water/William’s E Medium 50/50 v/v) were used and a quadratic model was for all compounds except ETU for which a cubic model gave a better fit. The samples were analyzed directly or after dilution with a mixture water/William’s E Medium (50/50 v/v) when appropriate. A control (calibration point at 5µM) was injected every 15 samples to check the calibration accuracy. The samples to be analyzed were culture media containing or not mancozeb and incubated or not with HepaRG cells.

### 2.7. Mn and Zn quantification by Inductively Coupled Plasma Mass Spectrometry (ICP-MS)

HepaRG cells were treated during 16 hours with maneb at 3ADI concentration, MnCl_2_ at 7µM or vehicle solvent (DMSO) as negative control. Then, cells were collected using trypsin (0.05%) before centrifugation. Cell pellets were washed twice in PBS then frozen at −80°C. Cells or purified mitochondria (see section 2.5) were desiccated overnight at 120°C. Dried tissues were weighed and mineralized by nitric acid solution in teflon PFA-lined digestion vessels. Acid digestion was carried out at 180°C using ultrapure concentrated HNO_3_ (69%) (Fisher Chemical Optima Grade) in a microwave oven device (Mars 6, CEM). The elements studied were ^55^Mn and ^66^Zn. ICP-MS analyses were carried out using an X-Series II from Thermo Scientific equipped with collision cell technology (Platform AEM2, Biochemical Laboratory, Rennes Hospital). The source of plasma was argon (purity degree > 99.99%). The collision/reaction cell used was pressurized with a mixture of helium (93%) and hydrogen (7%); argon and hydrogen were provided by Messer. Ultra-pure water was provided from Millipore Direct-Q 3 water station. Nitric acid solution was utilized at 69% (Fisher Chemical—Optima Grade). The rhodium was used as internal standard (Fisher Scientific). Calibration ranges were realized using a multi-element solution (SCP Science PlasmaCal). The performance was calibrated using multi-element solutions, tune F, and tune A (Thermo). The certified reference biological material was bovine liver ZC71001 obtained from NCS Testing Technology (Beijing, China).

### 2.8. RNA and protein expression studies

RNA purification and reverse transcription were performed using Nucleospin RNA (Fisher Scientific ref: 740955) and High capacity cDNA reverse transcription kit (Applied Biosystems). Quantitative PCR was performed with Sybr Green PCR Master Mix (Applied Biosystems) on ABI PRISM 7900HT instrument. The primer sequences are provided in **Supporting Information 1**.

For immunoblotting, culture medium was removed and cells were washed once with cold PBS and lysed in 50 mM HEPES pH 7.9, 150 mM, NaCl, 0.1 mM EDTA, 10% glycerol, 0.5% Tween 20 lysis buffer supplemented with protease inhibitors (EDTA-free, Roche). Cell lysates were briefly sonicated and total proteins of cell extracts were quantified using Biorad protein reagent assay. For detection of mitochondrial proteins, mitochondria were isolated from HepaRG cells after 16h of treatment with pesticides. Cells were lysed in H-medium (HEPES 5mM, D-mannitol 210mM, sucrose 70mM, pH 7.4). The lysate was centrifuged 5 minutes at 600g, 4°C. The supernatant (S1) was centrifuged 10 minutes at 4300g, 4°C. The supernatant (S2) was centrifuged 2h at 20,000g at 4°C. Resulting supernatant corresponded to cytosolic fraction. The pellet (C2) was then resuspended and centrifuged 5 minutes at 600g, 4°C. The supernatant collected (S3) was centrifuged 10 minutes at 4300g at 4°C. The pellet containing purified mitochondria was resuspended in a minimum volume of H-medium and frozen at −80°C.

Thirty µg of cytosolic or 10µg of mitochondrial proteins from each sample were separated on NuPAGE^®^ Novex^®^ Bis-Tris 4–12% gels kit (Invitrogen) and transferred to PVDF membranes (Trans-blot^®^ Turbo™ Transfer System, Biorad) prior to immunoblotting using the following primary antibodies: COX4 (F-8, sc-376731, Santa Cruz), GSTA1 (R-14, sc-100546 Santa Cruz), Anti-Cytochrome C (556433, BD Pharmingen), IRE1a (14C10, 3294 Cell Signalling), anti phospho S724-IRE1a (ab124945, Abcam), eIF2a (FL-315, sc-11386 Santa Cruz), anti-phospho Ser52-eIF2a (sc-12412, Santa Cruz) and HSC70 (B-6, sc-526-G Santa Cruz). Primary antibodies were detected using secondary antibodies coupled to horseradish peroxidase (Dako, Denmark), and detection of the immune complex was performed by chemiluminescent detection (Pierce™ ECL Substrate).

### 2.9. Data analysis

All data are expressed as mean ± standard error of the mean (SEM) from at least three independent experiments. Comparisons between groups were performed using one-way ANOVA followed by a post hoc Dunnett’s test or Bonferroni’s test when the normality test was positive, and Kruskal-Wallis followed by a post hoc Dunn’s test when data were not normally distributed. Graph Pad Prism 8 software (GraphPad Software, San Diego, CA, USA) was used for all statistical analyses) and the threshold for statistical significance was set to *p* < 0.05.

## 3. Results

### 3.1. Chronic and acute exposures to the pesticide mixture induce cytotoxicity in human hepatocytes

In order to study the effects of the mixture of the 7 pesticides on human hepatic cell fate, we performed chronic exposures of human progenitor HepaRG cells at %ADI, ADI and 3ADI pesticide concentrations during 30 days (**Figure 1**) as previously done for BM-MSCs (Hochane et al., 2017; Foucault et al., 2021).

**Figure 1.**
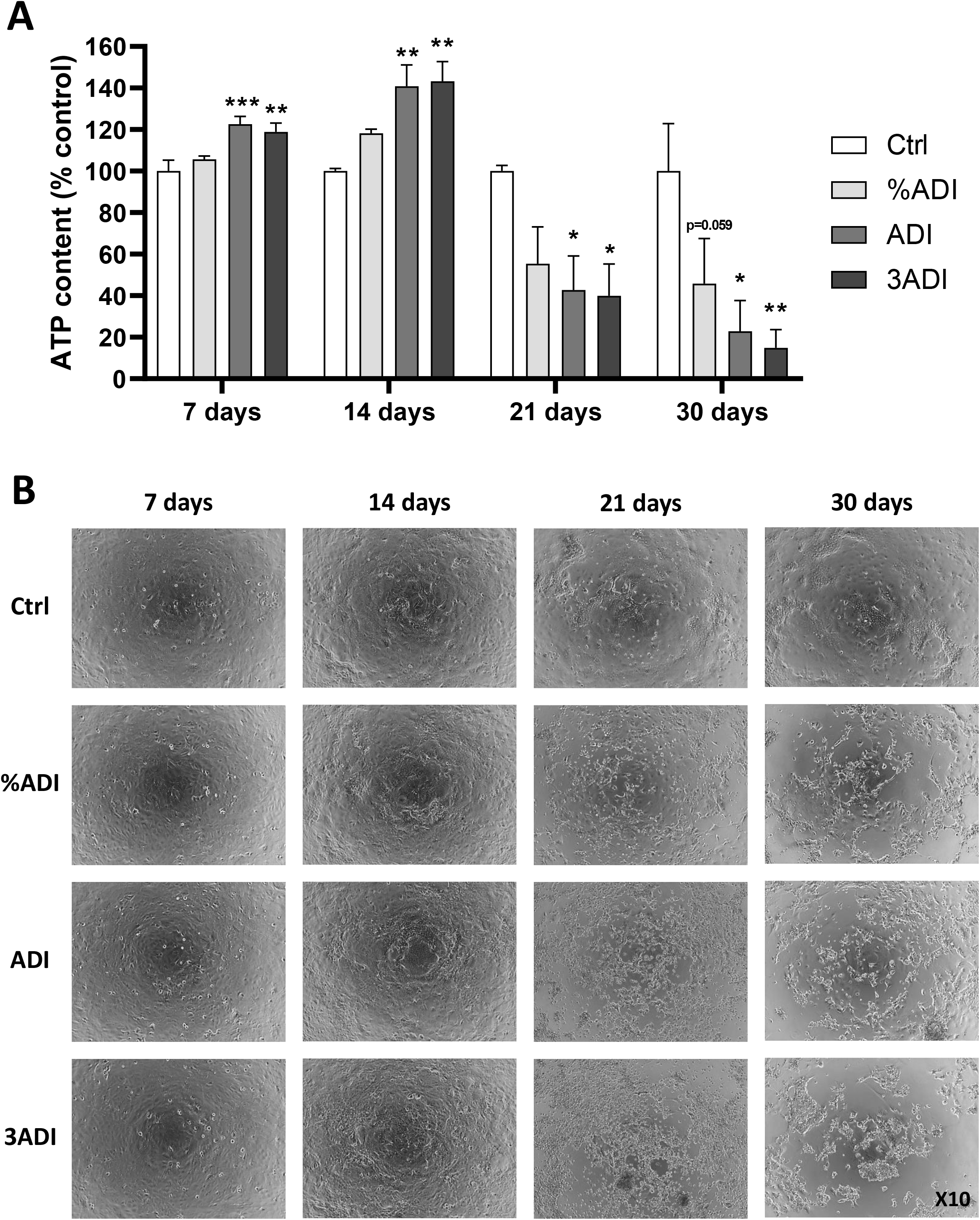
Chronic exposure of HepaRG cells to the pesticide mixture at low concentrations. **A**. Progenitor HepaRG cells were treated with the pesticide mixture during 4 weeks at different concentrations (%ADI, ADI or 3ADI). Cell viability was evaluated by measuring the intracellular ATP content after 7, 14, 21 and 30 days of exposure. Data were expressed as percentages of untreated control cells. Results are means ± SEM (N≥3), *p< 0.05, **p< 0.01 significantly different from control (Ctrl) using One-way ANOVA and Dunnett’s multiple comparisons test. **B**. Cell morphologies in phase contrast of control and pesticide-treated cultures.

After plating at low density, progenitor hepatoblast-like HepaRG cells actively proliferate during 7 to 10 days prior differentiation into hepatocyte and cholangiocyte cell lineages (Cerec et al., 2007; Vlach et al., 2019). During the first two weeks, the pesticides at the ADI and 3ADI concentrations induced a slight but statistically significant increase in ATP contents (**Figure 1A**) without any visible effect on the cell morphology (**Figure 1B**) suggesting that the phytosanitary compounds could stimulate the proliferation of progenitor HepaRG cells. In contrast, when the chronic treatment was prolonged for 2 more weeks during differentiation, the pesticides triggered a strong decreased in the ATP content in a concentration dependent manner (**Figure 1A**). At 21 days of treatment, floating cells were visible in culture wells incubated to pesticides and very few cells remained adherent at 30 days (**Figure 1B**). These data demonstrated that the pesticide mixture triggered a dose- and time-dependent cytotoxicity in differentiated HepaRG cells.

In order to determine whether the cytotoxicity of the pesticides observed after 2 weeks resulted from the chronic exposure of HepaRG cells during proliferation or occurred only when cells underwent differentiation, we performed acute exposures of differentiated HepaRG cells (**Figure 2**). The cells were first differentiated during 30 days in absence of pesticides, then subjected to a single exposure to the pesticide mixture. After 2 days, we observed a strong decrease in the ATP content of differentiated HepaRG cells in a dose-dependent manner. Significant effects were found with concentrations as low as ½ ADI but the overall reduction in cell viability reached 50% of the control value for the ADI and higher concentrations. These data demonstrated that the pesticide mixture induced an acute toxicity in hepatocytes *in vitro* after a single exposure.

**Figure 2.**
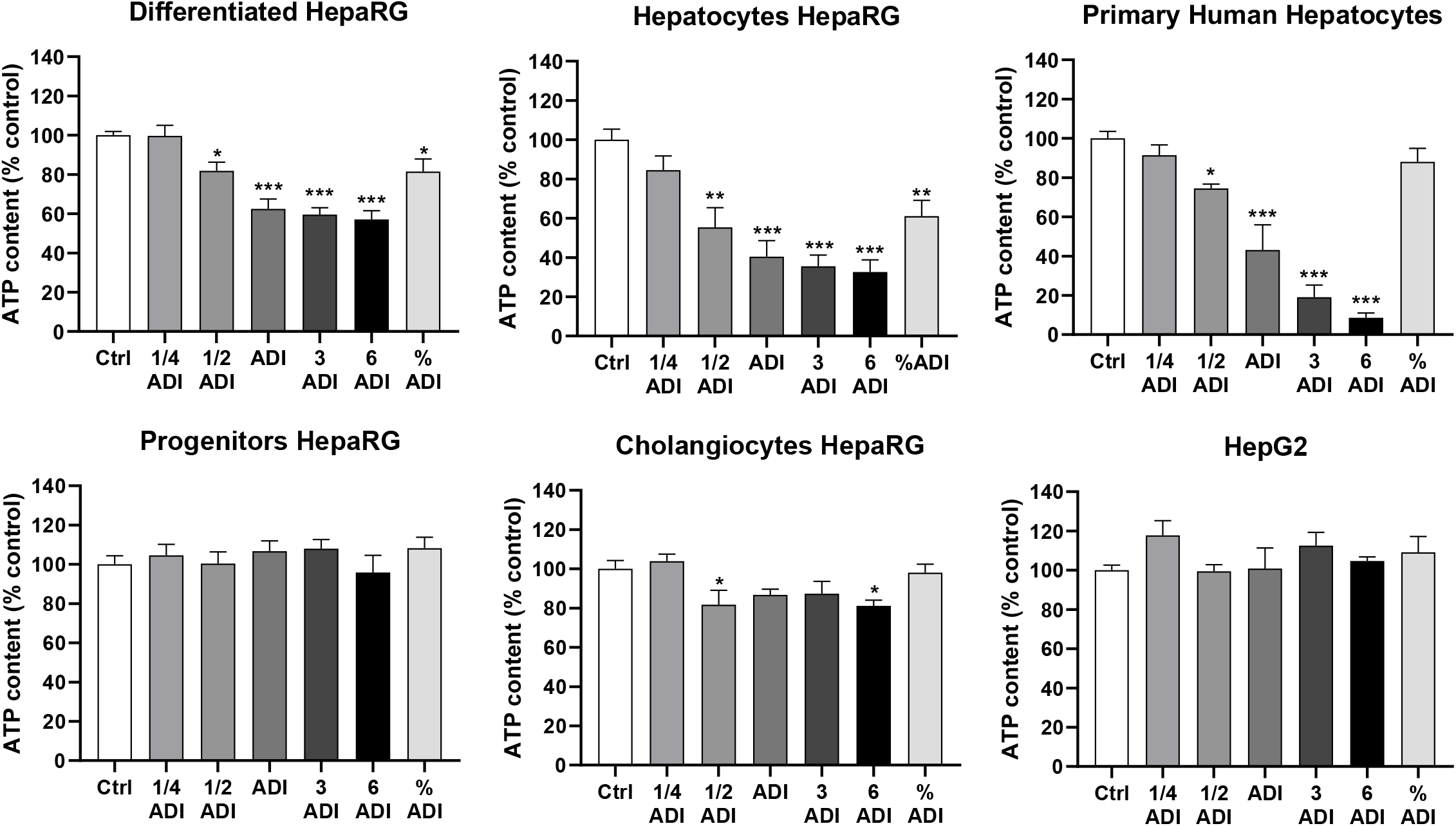
Acute exposure of hepatic cells to the pesticide mixture at low concentrations. Cell viabilities of differentiated HepaRG cells, human hepatocytes in primary culture and HepG2 hepatoma cells were measured by quantification of the intracellular ATP content after 48 h of exposure at different concentrations of the pesticide mixture (1/4ADI, 1/2ADI, ADI, 3ADI, 6ADI or %ADI). Data were expressed as % of untreated control cells. Results are means ± SEM (N≥3), *p< 0.05, **p< 0.01, ***p< 0.001 significantly different from control (Ctrl) using One-way ANOVA and Dunnett’s multiple comparisons test.

Differentiated HepaRG cultures contain both hepatocyte- and cholangiocyte-like cells producing a coculture *in vitro* model (Cerec et al., 2007). We next investigated whether the pesticides affected differently the viability of the 2 cell types by preparing enriched populations of both hepatocyte- and cholangiocyte-like HepaRG cells incubated in the same conditions (**Figure 2**). Interestingly, the pesticides triggered a stronger cytotoxicity in the hepatocyte-like cell population compared to that measured in the coculture while a mild decrease in ATP content was observed in the cholangiocyte cultures. The selective cytotoxicity towards hepatocytes was also visible with the morphological alterations in hepatocyte-like cell colonies while cholangiocytes remained unaffected (**Supporting Information 3A**).

To further confirm the toxicity of the pesticide cocktail, we used pure cultures of PHH incubated in the same conditions (**Figure 2**). The ATP content decreased in a dose-dependent manner with an extended loss in cell viability at 3 and 6ADI concentrations of pesticides. These data demonstrated the high cytotoxic effect of the pesticide cocktail towards differentiated hepatocytes. This conclusion was further supported by the absence of cytotoxicity in progenitor HepaRG cells, human HepG2 (**Figure 2**) and HuH7 hepatoma cells as well as primary rat liver epithelial SDVI cells, human macrophages and mesenchymal cells (**Supporting Information 3B-C**). In addition, we also used 2 transgenic HepaRG cell lines expressing the green fluorescent protein (GFP) under transcriptional control of hepatocyte-specific CYP2B6 and CYP3A4 gene promoters (Vlach et al., 2019) leading to GFP expression at high levels in hepatocyte-like HepaRG cells (**Supporting Information 3D**). Upon exposure to the pesticide mixture, the GFP positive hepatocyte colonies were strongly reduced in size and fluorescence intensities. Moreover, in transgenic HepaRG cells exposed to pesticides, the mRNA levels of the endogenous CYP3A4 and CYP2E1 were dramatically decreased compared to those found in untreated cells while the expression of cytokeratin 19 (CK19), which is expressed in cholangiocytes remained stable (**Supporting Information 3E**). Together, our data demonstrated that the pesticide cocktail triggered cytotoxicity specifically in hepatocytes.

### 3.2. Maneb and mancozeb induce oxidative stress and apoptosis in human hepatocytes

In order to determine whether one or several pesticides within the mixture were responsible for the toxicity, we exposed hepatocyte-like HepaRG cells to each pesticide individually in a wide range of concentrations (from ADI to 50xADI) for 48h and evaluated cell viability by measuring the ATP content (**Figure 3A**). Exposures to each pesticide have unveiled that only maneb and mancozeb were toxic. Moreover, the decreases in ATP content were similar with one or both DTCs and the mixture of all compounds (**Figure 2**) demonstrating the absence of additive effects of maneb and mancozeb and strongly suggesting that the toxicity observed with the mixture was triggered by these DTCs. To confirm this hypothesis, we exposed hepatocyte-like HepaRG cells to 7 different mixtures depleted in each pesticide individually and a mixture of 5 pesticides without maneb and mancozeb (**Figure 3B**). All mixtures depleted in one compound induced the same decrease in ATP contents while the cocktail of 5 pesticides without the two DTCs had no effect on cell viability confirming that maneb and mancozeb induced the toxicity of the mixture and evidencing for the first time that these DTCs trigger cell death in human hepatocytes at very low concentrations in the range of the ADI.

**Figure 3.**
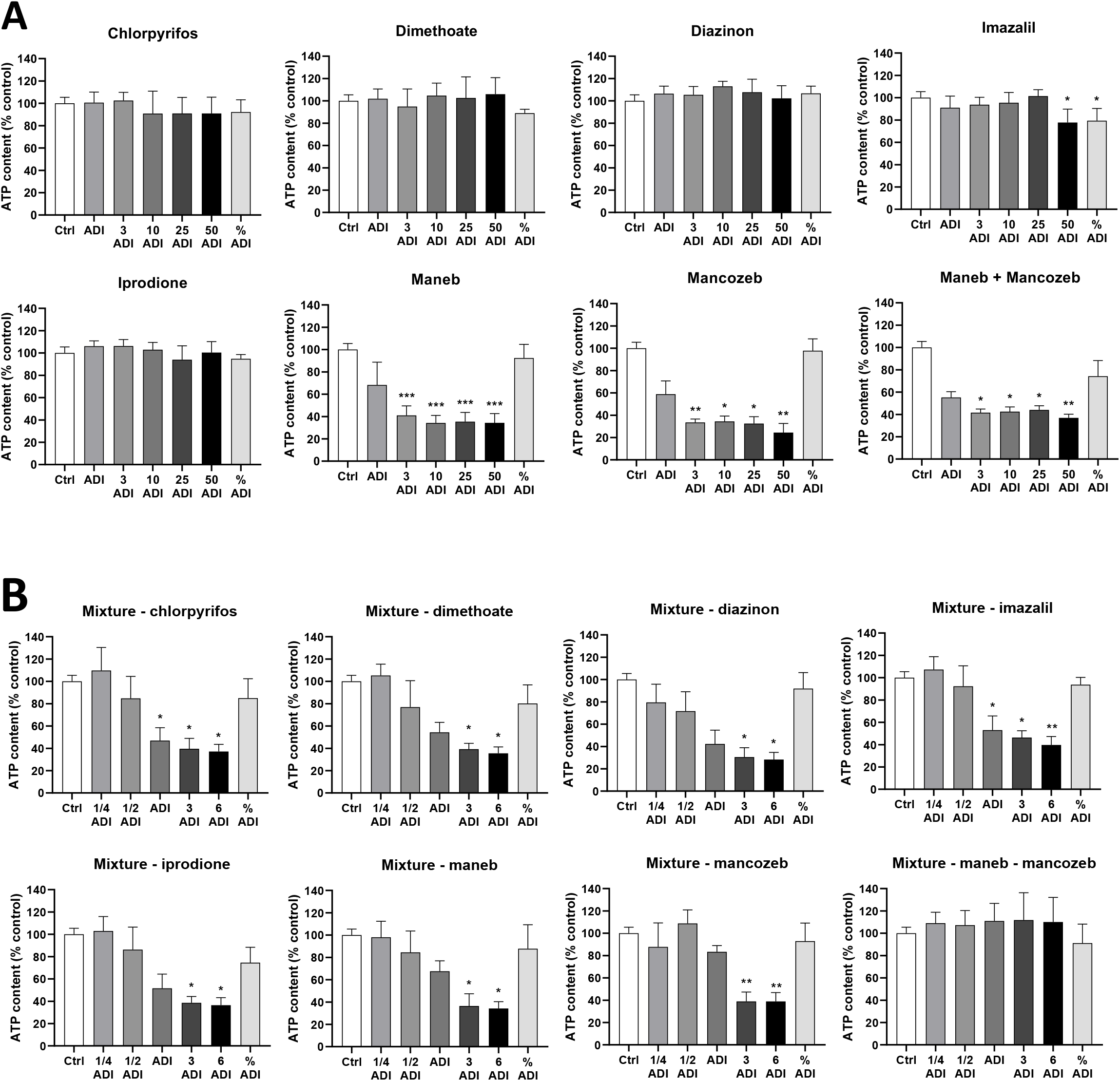
Hepatotoxicity of the pesticide mixture is triggered by maneb and mancozeb. **A**. Cell viability was evaluated by measuring intracellular ATP contents in hepatocyte-like HepaRG cells either in untreated control cultures (Ctrl) or exposed for 48h to each pesticide individually and the coincubation with maneb and mancozeb at different concentrations (%ADI, ADI, 3ADI, 10ADI, 25ADI, 50ADI). **B**. Cell viability (ATP content) in hepatocyte-like HepaRG cells incubated for 48 h to 7 pesticide mixtures depleted in each pesticide (6 compounds per mixture) and a cocktail of 5 pesticides without maneb and mancozeb at different concentrations (%ADI, 1/4ADI, 1/2ADI, ADI, 3ADI, 6ADI). Data were expressed as % of control. Results are means ± SEM (N≥3), *p< 0.05, **p< 0.01 significantly different from control (Ctrl) using One-way ANOVA + Dunnett’s or Kruskal Wallis + Dunn’s multiple comparisons tests.

We next studied the mechanism of cytotoxicity induced by the pesticide mixture and DTCs (**Figure 4**) postulating that maneb and mancozeb may induce apoptosis in hepatocytes since previous publications reported that mancozeb at high concentrations (10 to 300 μM) triggered apoptosis in human gastric, colon and hepatoma cells (Kumar et al., 2019; Dhaneshwar and Hardej, 2021; Lori et al., 2021). We measured caspase 3, 8 and 9 catalytic activities in total cell lysates of hepatocyte-like HepaRG cells exposed to the pesticide mixture or mancozeb alone (**Figure 4A**). The initiator caspase 9 and executioner caspase 3 activities were significantly increased in cells incubated with the mixture and mancozeb while the caspase 8 activity was unchanged suggesting the lack of cell death receptor activation. Importantly, when the antioxidant NAC was added prior to the pesticide mixture, the increase in caspase 3 and 9 activities was totally abrogated. Similarly, the pesticide mixture strongly induced the caspase 3 activities in human hepatocytes in primary culture (**Figure 4B**). The induction of apoptosis was further demonstrated by the cytochrome *c* release from mitochondria in both differentiated HepaRG cells and human hepatocytes (**Figure 4C**) and the increase in subG1 fraction of hepatocyte-like HepaRG cells (**Supporting Information 4A**). Together, these data demonstrated that the pesticide mixture and mancozeb induced caspase-dependent apoptosis in human hepatocytes through an intrinsic pathway and suggested the involvement of oxidative stress since NAC prevented the onset of cell death.

**Figure 4.**
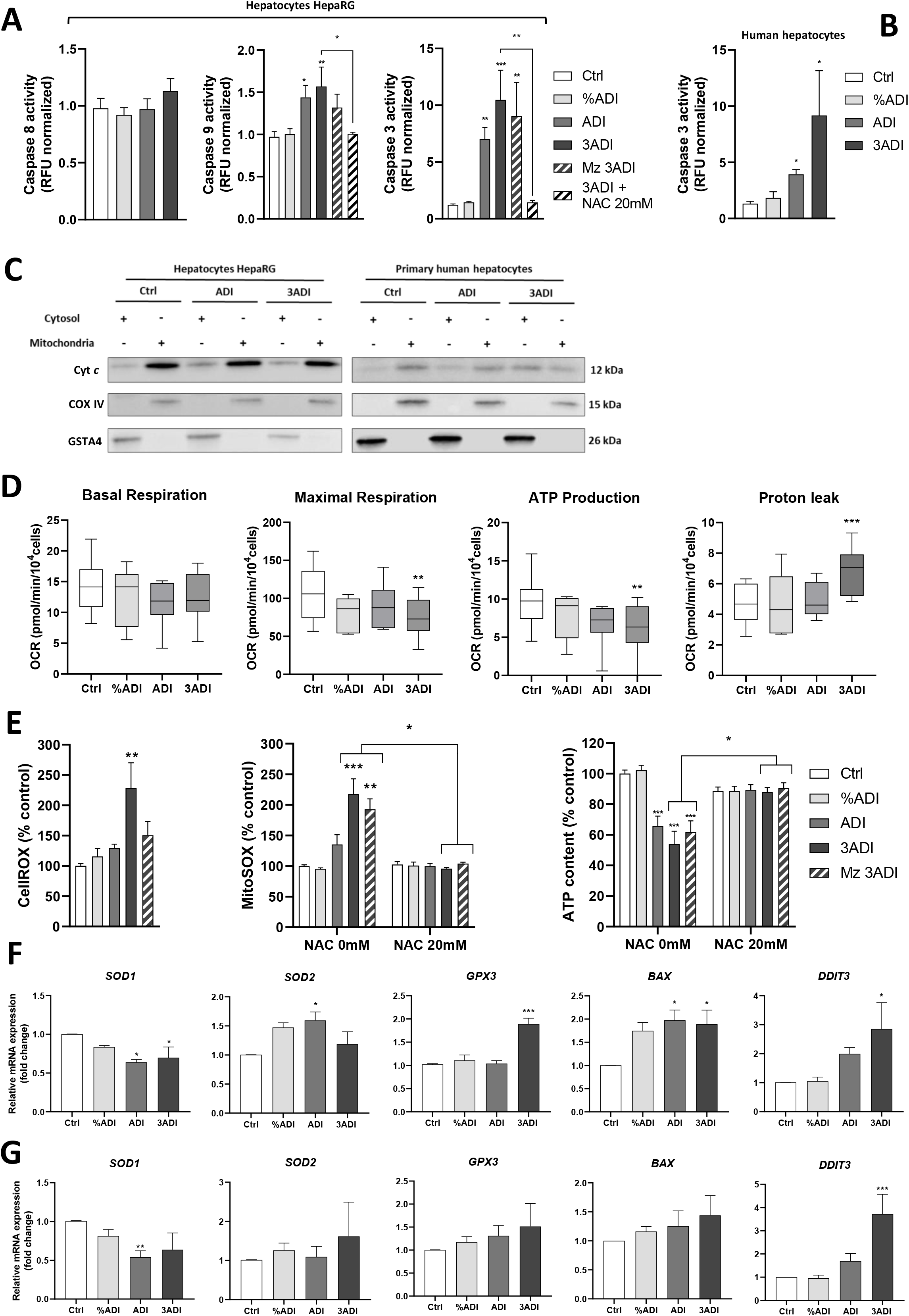
Pesticide mixture and mancozeb induce oxidative stress and apoptosis in human hepatocytes. **A**. Caspase 8, 9 and 3 activities were quantified using Ac-IETD-AMC, Ac-LEHD-AMC and Ac-DEVD-AMC fluorogenic substrates, respectively, in hepatocyte-like HepaRG cells exposed for 24h to the pesticide mixture at different concentrations (%ADI, ADI and 3ADI), in absence or presence of N-acetylcysteine (NAC, 20 mM), and mancozeb (Mz) at the 3ADI concentration. **B**. Ac-DEVD-AMC substrate caspase 3 activity quantified in human hepatocytes in primary culture exposed to the pesticide mixture for 48h at different concentrations (%ADI, ADI and 3ADI). Relative data were normalized to the basal values measured in untreated control cells arbitrarily set as 1 (Ctrl). Results are means ± SEM (N≥3), *p< 0.05, **p< 0.01, ***p< 0.001 significantly different from control (Ctrl). **C.** Detection of cytochrome *c* by western blot in cytosol and mitochondrial subcellular fractions of differentiated HepaRG cells and human hepatocytes in untreated control (Ctrl) conditions and exposed to the pesticide mixture for 24 h at the ADI and 3ADI concentrations. **D**. Parameters of mitochondrial respiration (basal and maximal respiration, ATP production and proton leak) were analyzed in differentiated HepaRG cells exposed for 16 h to the pesticide mixture (%ADI, ADI and 3ADI concentrations) using XF Cell Seahorse Mito Stress Test [Results are means ± SEM (N≥3), One-way ANOVA + Bonferroni’s multiple comparisons test or Kruskal Wallis + Dunn’s multiple comparisons test]. **E**. Oxidative stress was evaluated in hepatocyte-like HepaRG cells using CellROX and MitoSOX probes after 24 h exposure to pesticide mixture (%ADI, ADI or 3ADI) or mancozeb alone (Mz 3ADI). N-acetyl cysteine (NAC 20mM) was used to neutralize oxidative stress (Kruskal Wallis + Dunn’s multiple comparisons test). Cell viability of hepatocyte-like HepaRG cells was quantified by the relative intracellular ATP content after 24 h exposure to pesticide mixture (%ADI, ADI or 3ADI) or mancozeb alone (Mz 3ADI). **F-G**. Relative mRNA levels of *SOD1* and *2*, *GPX3*, *BAX* and *DDIT3* genes in hepatocyte-like HepaRG cells (**F**) and human hepatocytes (**G**) exposed for 24 h to the pesticide mixture (%ADI, ADI or 3ADI). Data were expressed as % of the untreated control culture (Ctrl). Results are means ± SEM (N≥3), *p< 0.05, **p< 0.01, ***p< 0.001, significantly different from control (Ctrl).

The effects of pesticides on mitochondrial function were next assessed in differentiated HepaRG cells. While basal respiration was not affected, maximal respiration and respiration linked to ATP production were significantly decreased upon exposure to the pesticide mixture at the 3ADI concentration (**Figure 4D**). In contrast, proton leak was increased by the same concentration. The activities of the five OXPHOS mitochondrial complexes were also studied in control culture and in cells exposed to maneb and no significant differences in enzymatic activities could be observed between untreated and pesticide-exposed cells (**Supporting information 4B**), thus suggesting an impairment of other metabolic pathways linked to OXPHOS.

The oxidative stress was evaluated by spectrofluorometry using the CellROX and MitoSOX probes, which are recommended to measure general oxidative stress and mitochondrial superoxide, respectively. While the CellROX indicator evidenced oxidative stress only at high concentrations of pesticides, relative superoxide levels detected with the MitoSOX probe were strongly enhanced in cells exposed to the pesticide mixture or mancozeb alone at ADI and 3ADI concentrations (**Figure 4E**). Importantly, NAC totally prevented pesticide-induced oxidative stress measured with MitoSOX probe and the cytotoxicity evaluated by the ATP content. In addition, no significant change in the relative ROS levels were observed in cholangiocyte-like HepaRG cells treated with the same concentrations of pesticides (**Supporting Information 4C**). Oxidative stress was further investigated in hepatocyte-like HepaRG cells (**Figure 4F**) and human hepatocytes (**Figure 4G**) by measuring the mRNA levels of genes known to be regulated by ROS and/or involved in antioxidant responses. Superoxide dismutase 1 (SOD1) mRNA levels were decreased while those of SOD2, glutathione peroxidase 3 (GPX3), BCL2-associated X apoptosis regulator (BAX) and DNA damage inducible transcript 3 (DDIT3, also known as CHOP) were up-regulated upon exposure to the pesticide mixture. The increase in DDIT3 transcripts suggested the induction of an endoplasmic reticulum (ER) stress, which was confirmed with the up-regulation of PERK and ERN1 mRNA levels and the increase in IRE1α phosphorylation and eIF1α protein expression (**Supporting Information 4D**). Considering the evidence of ROS overproduction and ER stress response in hepatocyte-like HepaRG cells incubated with the pesticides, a genotoxic effect was investigated using the γ-H2AX assay (**Supporting information 4E-F**). A strong induction in the percentage of γ-H2AX positive cells was detected and quantified in cells exposed to the pesticide mixture and aflatoxin B1 used as a positive control of genotoxicity. Altogether, these results indicated that the pesticide mixture and DTCs triggered a burst of ROS inducing ER stress, mitochondrial dysfunction, genotoxicity and apoptosis through an intrinsic caspase pathway.

### 3.3. Mancozeb is metabolized in hepatocytes to produce ethylene thiourea

Since cell death induced by DTCs occurred only in differentiated hepatocyte-like HepaRG cells and human hepatocytes, we investigated the catabolism of the mancozeb in these metabolically competent cells expressing phase I and II xenobiotic-metabolizing enzymes (XME) (Aninat et al., 2006; Kanebratt and Anderson, 2008), and in cholangiocyte-like and progenitor HepaRG cells, which do not express these XME and do not undergo apoptosis upon exposure to DTCs.

Detection of mancozeb was assessed by HPLC coupled to UV detector (HPLC-UV) (**Figure 5**) and mancozeb’s metabolites were identified by LC-MS/MS (**Table 1**, **Supporting Information 2**). Mancozeb added to culture media at the concentration of 100 µM was detected by HPLC-UV at 285nm with a retention time of 10.75 min (**Figure 5**). This peak was confirmed to be mancozeb by LC-MS/MS (**Table 1**, **Supporting Information 2**). In addition, in media supplemented with mancozeb, a second peak was detected by HPLC at 247nm at a retention time of 2.7 min, very similar retention time to that of ethylene thiourea (ETU) analytical standard (**Figure 5**). Using LC-MS/MS (**Table 1**), we confirmed the presence of ∼21 μM of ETU in culture media without incubation with HepaRG cells strongly suggesting the presence of this degradation product in the mancozeb stock solution. In culture media of hepatocyte-like HepaRG cells, the amounts of mancozeb progressively decreased during a 24h incubation time while those of ETU concomitantly increased (**Figure 5**, **Table 1**). In contrast, the peaks of mancozeb and ETU detected by HPLC in culture media of cholangiocyte-like and progenitor HepaRG cells were not quantitatively modified during the 24h incubation with these cells (**Figure 5**). These data demonstrated that mancozeb is catabolized into its main ETU metabolite in hepatocyte-like HepaRG cells and that non-hepatocytic cells do not metabolize mancozeb.

**Figure 5.**
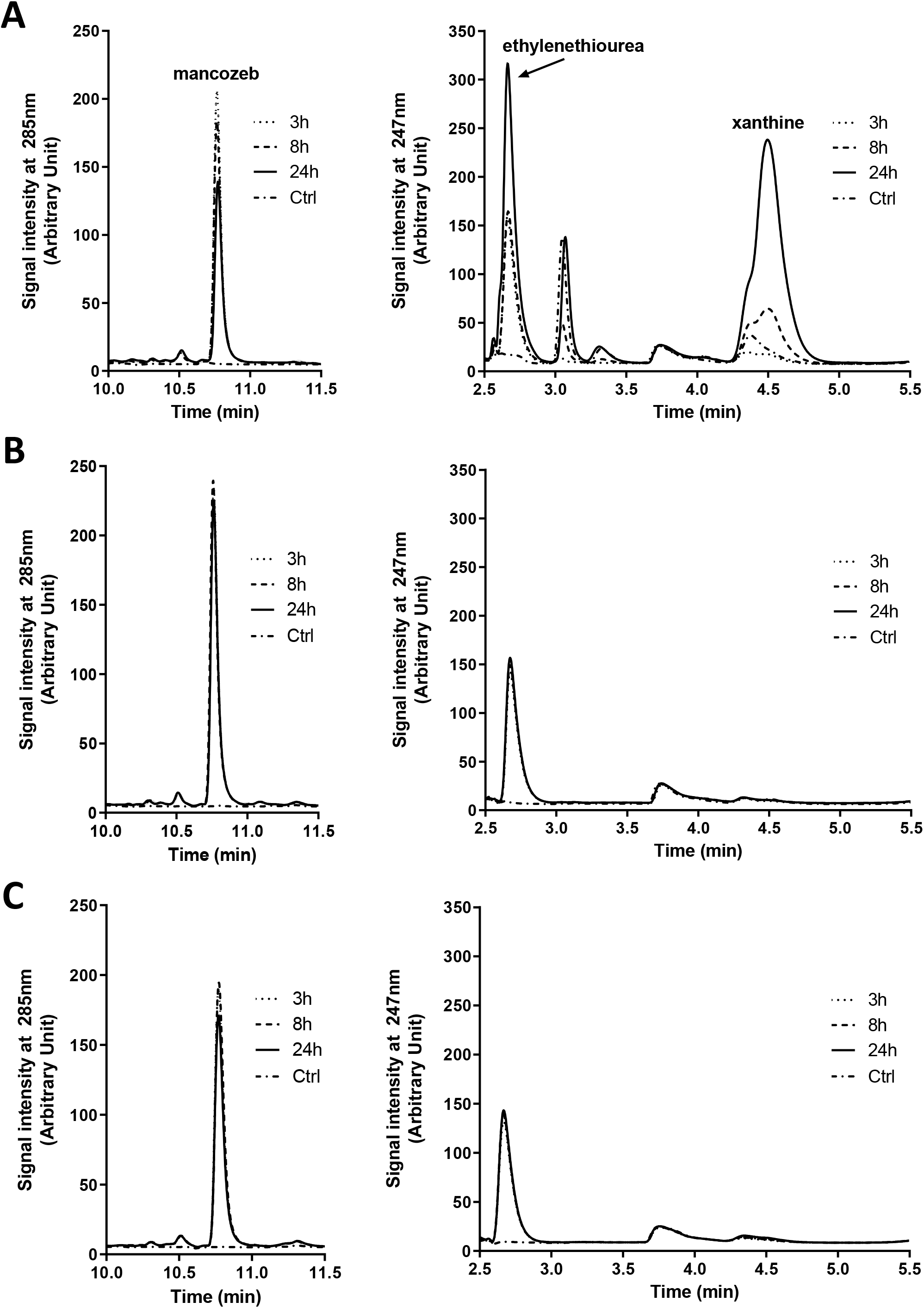
Hepatocyte-like HepaRG cells metabolize mancozeb. Representative UV-HPLC chromatograms of culture media without mancozeb (untreated control cells at 24 hours (h), Ctrl) or containing mancozeb (100µM) and incubated for 3, 8 and 24 h with **A**) hepatocyte-like, **B**) cholangiocyte-like and **C**) progenitor HepaRG cells. Mancozeb, ethylenethiourea (ETU) and xanthine peaks are indicated in panel A.

**Table 1:** Quantification of mancozeb, ethylene thiourea, hypoxanthine, xanthine and uric acid by LC-MS/MS (expressed in μM ±SD) and lactate dehydrogenase enzymatic activity(LDH, expressed in mU/mL) in culture media of hepatocyte-like HepaRG cells after 3, 8 and 24 hours of incubation with mancozeb at 100 μM. Statistics: p<0.05 *a* significantly different from 3h, *b* significantly different from 8h, * significantly different from untreated control (DMSO), # significantly different from Mz 100μM.

|  | Hours | Mancozeb | Ethylene thiourea | Hypoxanthine | Xanthine | Uric acid | LDH (mU/mL) |
| --- | --- | --- | --- | --- | --- | --- | --- |
| **DMSO** | 3 | <0.05 | <0.1 | 1.70 ± 0.40 | 1.42± 0.30 | 0.64 ± 0.42 | 2.15 ± 1.68 |
|  | 8 | <0.05 | <0.1 | 0.48 ± 0.10 | 2.08± 0.82 | 2.21± 2.35 | 3.92 ± 0.89 |
|  | 24 | <0.05 | <0.1 | 0.75 ± 0.35 | 2.69 ± 3.36 | 10.34 ± 8.21 | 8.19 ± 3.66 |
| **Mancozeb (100μM)** | 3 | 118.28 ± 6.82 | 21.40 ± 0.65 | 0.62 ± 0.22 | 0.75 ± 0.16 | 0.32 ± 0.16 | 2.19 ± 0.86 |
|  | 8 | 104.79 ± 4.52 <sup>a</sup> | 20.73 ± 0.71 | 1.28 ± 0.93 <sup>a</sup> | 2.29± 0.69 <sup>a</sup> | 2.48 ± 1.98 | 4.10 ± 0.71 |
|  | 24 | 35.44 ± 2.91 <sup>a,b</sup> | 64.58 ± 2.58 <sup>a,b</sup> | 5.15 ± 3.29 <sup>a</sup> | 10.36 ± 3.63 <sup>a</sup> | 9.47 ± 5.49 | 10.10 ± 1.96 |
| **MnCl<sub>2</sub> (100μM)** | 3 | <0.05 | <0.1 | 0.99 ± 0.45 | 1.00 ± 0.18 | 0.27 ± 0.14 | 1.97 ± 0.61 |
|  | 8 | <0.05 | <0.1 | 2.97 ± 2.78 | 3.92 ± 1.08 | 1.59 ± 1.18 | 4.46 ± 0.94 |
|  | 24 | <0.05 | <0.1 | 6.96 ± 3.39 <sup>a,*</sup> | 13.66 ± 0.11 <sup>a,*</sup> | 9.22 ± 1.71 | 14.37 ± 1.96 <sup>*</sup> |

In addition to the peaks of mancozeb and ETU, we also detected by HPLC-UV two other compounds eluting with retention times of 3.1 and 4.5 min only in culture media of hepatocyte-like HepaRG cells (**Figure 5A**). The peak at 3.1 min was not detected after 3 hours of incubation with mancozeb in hepatocyte-like HepaRG cells. However, the area of this peak increased with the incubation times and was similar in both untreated and treated cultures at 24h demonstrating that it corresponded to an endogenous hepatocyte-like HepaRG metabolite unrelated to mancozeb metabolism. This hypothesis was further confirmed by the absence of this peak in media of cholangiocyte-like HepaRG cells (**Figure 5B**) cultured in the same medium than hepatocyte-like HepaRG cells. Besides the formation of its well-documented ETU metabolite, the catabolism of mancozeb involves the production of at least 12 other metabolites (Engst and Schnaak, 1970; World Health Organization 1988; IRNS 2010), most of them known to be unstable compounds difficult to detect (**Supporting Information 2**). In a detailed LC-MS/MS analytical study, we investigated the presence of these metabolites in the culture media of hepatocyte-like HepaRG cells in order to identify the unknown compound eluting at 4.5 min in HPLC chromatograms (**Supporting Information 4**). We found that all known mancozeb metabolites were either undetectable or present at concentrations below the quantification limits (**Supporting Information 2**). By LC-MS/MS analysis, we found that the unknown compound eluting at 4.5 min in HPLC-UV had a molecular mass of 152 g/mol. Using the MASS BANK database, we identified two putative candidates, xanthine and oxipurinol, two compounds of the purine metabolic pathway. The LC-MS/MS analysis of commercial standards of these two compounds demonstrated that the unknown compound was xanthine (**Supporting Information 2**, **Table 1**). Although xanthine was detected in the culture media of control cells unexposed to mancozeb (∼2.7 μM at 24h of culture), its levels were significantly increased up to 10.4 μM in media of hepatocyte-like HepaRG cells incubated with 100 μM mancozeb (**Table 1**). We next analyzed hypoxanthine and uric acid, two other main compounds of the purine metabolic pathway. We found that hypoxanthine was also produced by hepatocyte-like HepaRG cells incubated with mancozeb but not uric acid. Quantification of the lactate dehydrogenase (LDH) activities showed no significant differences between culture media of cells exposed or not to mancozeb (**Table 1**) indicating that the increase in hypoxanthine and xanthine levels upon mancozeb treatment was due to an active secretion rather than a passive leakage from cells resulting from loss of plasma membrane integrity in mancozeb-treated cells. Since hypoxanthine and xanthine have not been identified as metabolites of DTCs, we hypothesized that their efflux was a stress response resulting from mancozeb exposure.

### 3.4. Exposure to DTCs leads to Mn overload and Zn depletion and apoptosis in hepatocytes

In order to determine if the production of DTC metabolites by hepatocyte-like HepaRG cells was responsible for DTC cytotoxicity, we exposed these cells to ETU as well as ethylene urea (EU) and ethylene diamine (EDA) and the combination of the three metabolites at concentrations ranging from 2 to 20 μM corresponding to mancozeb at ADI to 10xADI concentrations (**Figure 6A**). Exposure to these molecules had no significant effects on cell viability demonstrating that these metabolites were not involved in the cytotoxicity of DTCs towards hepatocytes.

**Figure 6.**
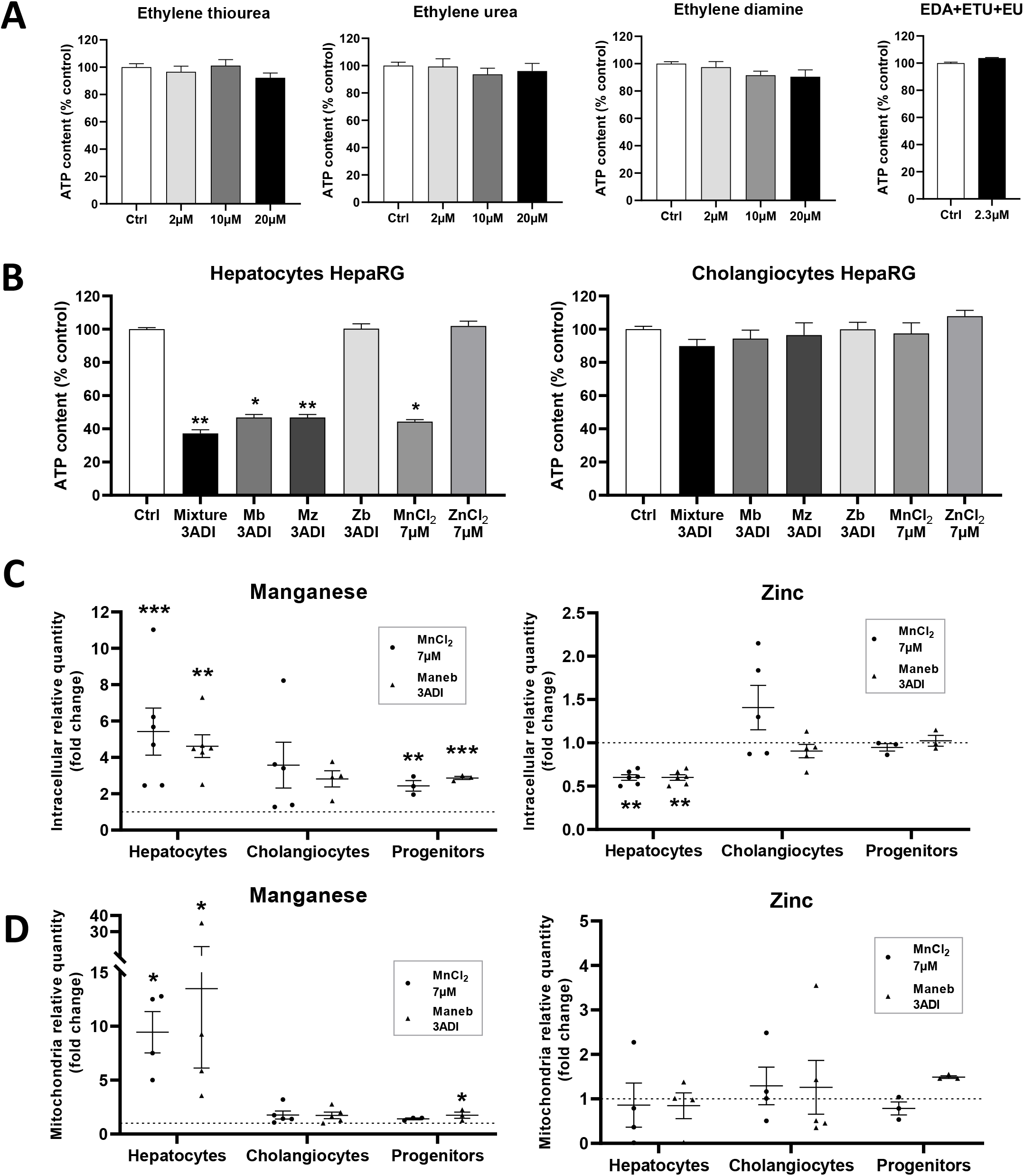
Exposure to maneb induces Mn overload and Zn depletion triggering cytotoxicity in hepatocytes. **A**. Effects of DTCs metabolites ethylene thiourea (ETU), ethylene urea (EU) and ethylene diamine (EDA) metabolites and the combination of the three molecules (EDA+ETU+EU) at 2, 10 and 20 μM on cell viability (relative ATP contents) in hepatocyte-like HepaRG cell after 48h of exposure. **B**. Cell viability (relative ATP contents) in hepatocyte- and cholangiocyte-like HepaRG cell cultures after 48h treatments with the pesticide mixture (mixture), DTCs maneb (Mb), mancozeb (Mz), zineb (Zb) at 3ADI concentration, MnCl_2_ and ZnCl_2_ at 7µM. Relative ATP contents are expressed as % of untreated control cultures (Ctrl). Results are means ± SEM (N≥3), *p< 0.05, **p< 0.01 significantly different from control (Ctrl) with Kruskal Wallis + Dunn’s multiple comparisons test. **C.** Relative intracellular (**C**) and mitochondrial (**D**) Mn and Zn contents in progenitor, hepatocyte- and cholangiocyte-like HepaRG cells measured by ICP/MS after 16h of exposures to maneb (3ADI) and MnCl_2_ at 7µM. Data are expressed as fold change in metal contents between unexposed control cells arbitrary set as 1 (dotted line) and treated cells because of the large differences in metal contents between cell types and intracellular versus mitochondrial amounts. Absolute quantities of both Mn and Zn are presented in ng/well in the Supporting Information 5. Results are mean ± SEM (N≥3), *p< 0.05, **p< 0.01 significantly different from control (Ctrl) One-way ANOVA + Dunnett’s multiple comparisons test or Kruskal-Wallis + Dunn’s multiple test

DTCs represent a large group of synthetic compounds that share a similar chemical structure (R_2_NCS_2_R) (Ajiboye et al., 2022) (**Supporting Information 2**). Several of these DTCs, including maneb, zineb and mancozeb, are salts containing Mn and Zn. While maneb and zineb contain only Mn and Zn, respectively, mancozeb is a combination of both maneb and zineb. We hypothesized that catabolism of DTCs in hepatocytes could lead to the release of Mn and/or Zn, present in their chemical structure. To address this hypothesis, hepatocyte-like HepaRG cells were exposed for 48h to maneb, mancozeb and zineb at the 3ADI concentration, as well as MnCl_2_ and ZnCl_2_ at 7µM corresponding to the concentration of Mn and Zn brought by exposure to DTCs at 3ADI (**Figure 6B**).

Quantification of the relative ATP contents in hepatocyte-like HepaRG cells demonstrated that exposure to maneb, mancozeb and MnCl_2_ triggered similar toxicities while zineb and ZnCl_2_ had no significant impact of cell viability (**Figure 6B**). The similar experiment performed using cholangiocyte-like cells showed no effect on cell viability with any of these treatments. These data strongly suggested that cell death of hepatocyte-like HepaRG cells exposed to maneb and mancozeb resulted from the release of Mn during their catabolism. We next studied the effects of exposures to maneb and MnCl_2_ on the intracellular contents in Mn and Zn measured by ICP/MS in progenitor, cholangiocyte- and hepatocyte-like HepaRG cells (**Figure 6C**-**D**, **Supporting Information 5**). Hepatocytes exposed to maneb and MnCl_2_ showed strong intracellular and mitochondrial accumulations of Mn with contents significantly higher than in untreated cells. Intracellular amounts of Mn were also increased in cholangiocytes and progenitor cells but in much lower extents compared those of hepatocyte-like cells. Importantly, significant decreases in intracellular Zn levels were also observed upon incubations with MnCl_2_ and maneb only in hepatocyte-like cells. Together, these data supported the conclusion that Mn-containing DTCs induced cell death in metabolically competent hepatocytes capable of metabolizing these fungicides to produce innocuous metabolites, such as ETU, and to release Mn accumulating in cells and altering Zn content.

Since the treatment of hepatocyte-like HepaRG cells by MnCl_2_ induced cytotoxicity, we used this experimental condition to determine whether the production of hypoxanthine and xanthine previously observed with mancozeb occurred in MnCl_2_-treated cells (**Figure 5**, **Table 1**). We found that exposure of differentiated HepaRG cells to MnCl_2_ also enhanced the secretion of both xanthine and hypoxanthine (**Table 1**) demonstrating that these compounds are not metabolites resulting from DTC catabolism and suggesting that they may be produced in response to the alteration of the intracellular Mn/Zn content and/or oxidative stress.

To further study the role of the Zn depletion in DTC- and Mn-dependent toxicity, hepatocyte-like HepaRG cells were exposed to the pesticide mixture, maneb, mancozeb and MnCl_2_ in absence or presence of a 10-fold excess of ZnCl_2_ (70μM) and oxidative stress, relative ATP content, caspase 3 and 9 activities and cellular Mn and Zn amounts were measured (**Figure 7**). First, incubation of differentiated HepaRG cells with MnCl_2_ led to an increase in relative mitochondrial superoxide levels measured with MitoSOX probe at similar levels to those observed in cultures exposed to maneb or mancozeb while fluorescence measured with the CellROX probe was increased with pesticide mixture and mancozeb only (**Figure 7A**). In addition, co-treatment with 70µM ZnCl_2_ prevented the oxidative stress measured with MitoSOX probe (**Figure 7A**), the decrease in ATP content (**Figure7B**) and the onset of apoptosis (**Figure 7C**) in cultures exposed to DTCs and MnCl_2_. Determination of cellular Mn and Zn contents in untreated control cells and cultures exposed to MnCl_2_ and maneb in presence of 70μM of ZnCl_2_ (**Figure 7D**, **Supporting Information 5**) demonstrated that the 10-fold excess in Zn partially but significantly reduced the intracellular and mitochondrial Mn overload and prevented Zn depletion induced by maneb and MnCl_2_. Unexpectedly, while the excess in Zn increased the intracellular Zn content in absence of maneb and MnCl_2_, it also reduced the mitochondrial contents in Zn in all culture conditions. These results demonstrated that partial restoration of Mn and Zn homeostasis disrupted by Mn-containing DTCs was sufficient to prevent oxidative stress and cytotoxicity in hepatocytes.

**Figure 7.**
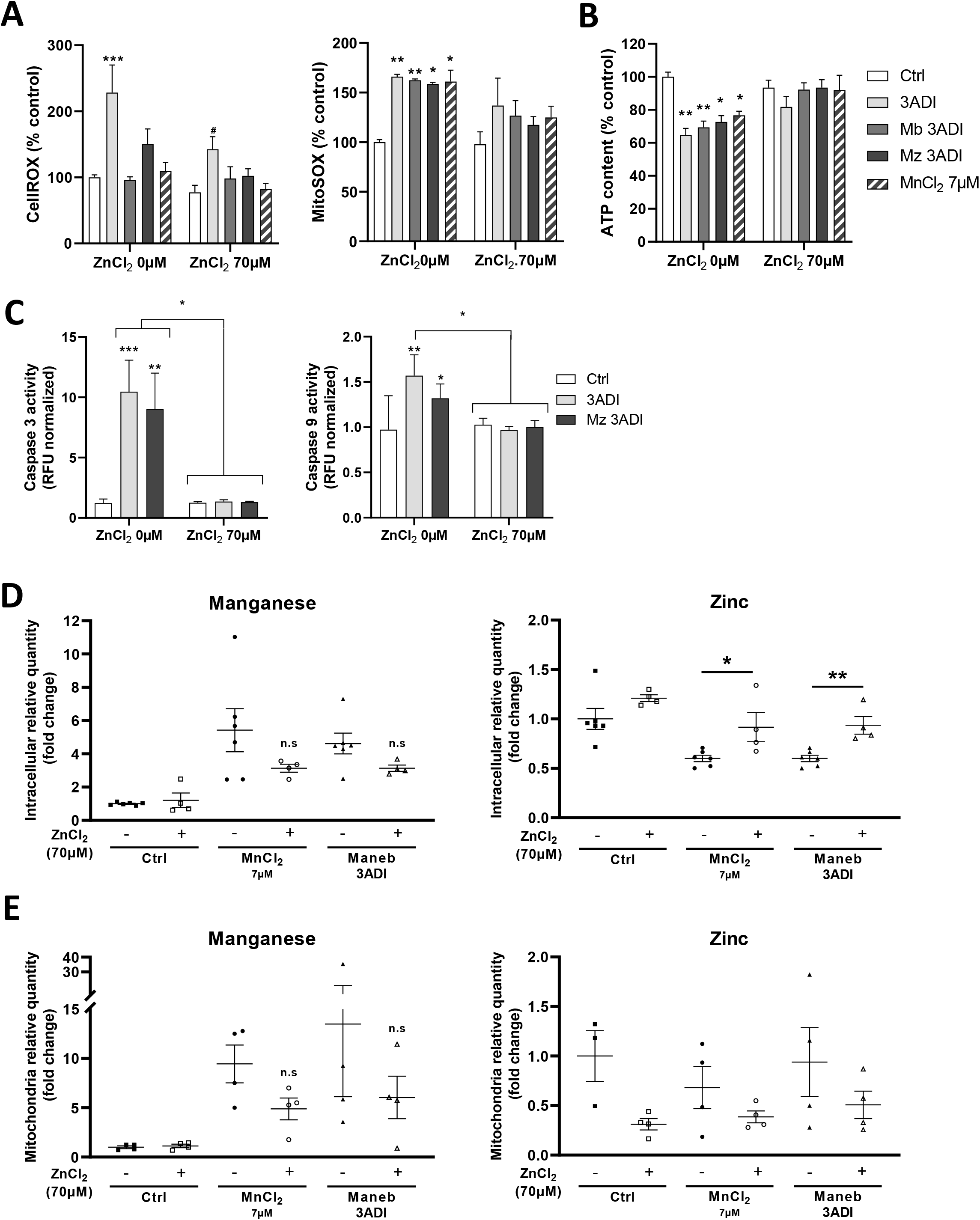
Excess of Zn prevents oxidative stress and cytotoxicity induced by DTCs in hepatocytes. **A**. Relative oxidative stress levels evaluated in hepatocyte-like HepaRG cells using CellROX and MitoSOX probes and, **B**. relative ATP contents in hepatocyte-like HepaRG cells after 24h of exposure to the pesticide mixture (3ADI), DTCs maneb (Mb 3ADI) and mancozeb (Mz 3ADI), and MnCl_2_ at 7µM in absence or presence of a 10-fold-excess in ZnCl_2_ at 70µM. For oxidative stress, statistics are: One-way ANOVA + Sidak’s multiple comparisons test or Kruskal-Wallis + Dunn’s multiple comparisons test, *p< 0.05, **p< 0.01, significantly different from untreated control cultures (Ctrl), #p< 0.05, significantly different from cultures exposed for 24h to the pesticide mixture, maneb, mancozeb and MnCl_2_ in absence of a 10-fold-excess in ZnCl_2_ at 70µM. For ATP contents, results are means ± SEM (N≥3), statistics are: Kruskal-Wallis + Dunn’s multiple comparisons test, *p< 0.05, **p< 0.01, significantly different from untreated control (Ctrl). **C**. Relative caspase 3 and 9 activities quantified using Ac-DEVD-AMC and Ac-LEHD-AMC florigenic substrates, respectively, after exposure to the pesticide mixture (3ADI) and mancozeb (Mz 3ADI) in absence or presence of a 10-fold-excess in ZnCl_2_ at 70µM. Kruskal-Wallis + Dunn’s multiple comparisons test. Results are means ± SEM (N≥3), *p< 0.05, **p< 0.01, ***p< 0.001, significantly different from untreated control cultures (Ctrl) and from cells co-exposed to excess of Zn. **D-E.** Relative intracellular (**D**) and mitochondria (**E**) Mn and Zn contents measured by ICP/MS after 16h exposures of hepatocytes HepaRG to maneb (3ADI) and MnCl_2_ at 7µM in absence (-) and presence (+) of a 10-fold excess of ZnCl_2_ (70µM). Data are expressed as fold change of untreated control cells (Ctrl). Results are means ± SEM (N≥3).

## 4. Discussion

The initial goal of this work was to study the effects of a mixture of 7 pesticides (chlorpyrifos-ethyl, dimethoate, diazinon, iprodione, imazalil, maneb, mancozeb) often detected in food samples (EFSA, 2017) at low concentrations in human hepatocytes *in vitro*. We evidenced that the pesticide mixture induced a strong cytotoxicity in hepatocytes-like HepaRG cells and human hepatocytes following chronic and acute exposures. The pesticides triggered the production of ROS, an ER stress response and DNA damages, which led to cell death by intrinsic apoptosis involving caspases 3 and 9. Unexpectedly, the individual evaluation of each pesticide demonstrated that the toxicity was carried by the Mn-containing DTCs maneb and mancozeb as the mixture of the five other pesticides without these compounds did not affect viability, and that the exposure to maneb or mancozeb alone recapitulated the effect of the mixture.

DTCs are broad-spectrum fungicides reacting with amino acids and enzymes. They are used on many fruits, vegetables, nuts, and field crops and spread on potatoes, corn, sorghum, tomatoes, apples, pears, grapes, onions and cereal grains to protect against fungal diseases. DTCs have been employed for decades with the examples of maneb and mancozeb registered on the market since 1948 and 1961, respectively. Nearly 200,000 tons of maneb are spread every year in the world and the overall amounts of mancozeb is estimated to several millions (Gullino et al., 2010; National Toxicology Program, 2014). Maneb was banned in 2017 in E.U. and but it is still used worldwide in large amounts. In 2018, the Agence Nationale de Sécurité Sanitaire de l’Alimentation, de l’Environnement et du Travail (ANSES) recommended to the European Commission the non-renewal of the mancozeb approval given risks for human health and the high levels of exposure resulting of its intensive use (ANSES, 2018). EU has adopted the non-renewal of mancozeb (Official Journal of the European Union, 2020), which was withdrawn in 2021 with a grace period that expired in January 2022.

Despite available data regarding DTC toxicity *in vivo* and *in vitro*, there are still ongoing investigations to decipher their mechanisms of action. Maneb and mancozeb show low acute toxicity *in vivo* (median lethal dose/LD_50_ >1g/kg), but adverse effects have been described in rodents upon chronic exposure (5 to 10 mg/kg b.wt./d) with neurotoxicity and hyperthyroidism (World Health Organization 1988; IRNS, 2010). Chronic occupational or experimental exposures to maneb induce Parkinson-like syndromes in humans and animals (Ferraz et al., 1988; Thiruchelvam et al., 2002; Anderson et al., 2021). Their main metabolite ETU also shows low toxicity (median lethal dose/LD_50_ >1.5g/kg b.wt.) in rodents. However, ETU is a structural analog of thionamides, which inhibits the thyroid peroxidase, reduces the synthesis of triiodothyronine (T_3_) and thyroxin (T_4_) and induces thyroid hypertrophy (Panganiban et al., 2004; Goldner et al.? 2010), defining an endocrine disrupting effect. ETU also alters the proliferation/survival of murine hematopoietic cell lineages with the development of pancytopenia after long-term exposure (Porreca et al., 2016). In addition, reports also suggest that occupational exposure to maneb and mancozeb induces alterations in lymphocyte and monocyte blood counts (Corsini et al., 2005; Colosio et al., 2007).

The *in vivo* hepatotoxicity of DTCs is less described with, however, studies demonstrating that mancozeb at high doses, 313.6 mg/kg b.wt./d, 3 times a week for 6 weeks (Sakr, 2007) and 750 mg/kg b.wt./d for 10 weeks (Hashem et al., 2018) induce liver damages with disruption of the normal histology, necrosis and mononuclear cell infiltration, depletion in glutathione, lipid peroxidation, genotoxicity and induction of inflammation in rats. Interestingly, antioxidant molecules such as Nigella sativa oil (Hashem et al., 2018) and curcumin (Saber et al., 2019) reduce the hepatotoxicity induced by mancozeb. Demonstration has also been brought that ETU (125mg/kg, single dose) induces hepatic morphological and histological abnormalities in newborns from female rats exposed during gestation, with strong remodeling of the hepatic trabeculae and severe hepatic megakaryocytosis (Lemos et al., 2012).

*In vitro* studies have described mitochondrial-mediated cytotoxicity and apoptosis of maneb and/or mancozeb in various cell types including rat mesencephalic primary embryonic cells (Domico et al., 2007), SK-N-AS neuroblastoma cells (Anderson et al., 2018), AGS gastric carcinoma cells (Kumar et al., 2019), HepG2 hepatoma cells (Lori et al., 2021) and HT-29 and Caco2 colon cells (Dhaneshwar et al., 2021, Hoffman and Hardej, 2012; Hoffman et al., 2016) through production of reactive oxygen species (ROS). In these reports, the cells were exposed to a wide range of concentrations of maneb and/or mancozeb from 1 to 500 μM, and the cytotoxic effects were observed at high concentrations from 10 to 100μM, except for rat mesencephalic primary embryonic cells in which ROS production was detected at 3μM (Domico et al., 2007).

To the best of our knowledge, our study is the first report describing induction of oxidative stress and apoptosis by maneb and mancozeb in differentiated hepatocyte-like HepaRG cells and primary human hepatocytes. In these cell models, the adverse effects occurred at concentrations as low as ½ADI (1.13 μM) in acute exposures and at hNDI (0.73 μM) in chronic treatments demonstrating that human hepatocytes are particularly sensitive to maneb- and mancozeb-induced toxicity. Interestingly, cytotoxic effects at such low concentrations were not found in other hepatoma cell lines and non-hepatic cells, which do not express phase I and II XMEs enabling pesticide metabolism. In this report, we showed that human hepatocytes metabolize mancozeb to produce ETU, which correlated with intracellular accumulation in Mn and depletion in Zn triggering ROS overproduction, xanthine/hypoxanthine release, ER stress and apoptosis.

It has been previously shown that maneb and mancozeb alter metal contents in HT-29 and Caco2 colon cells (Hoffman and Hardej, 2012; Hoffman et al., 2016). In these studies, the authors demonstrated that maneb and mancozeb cause Mn overload but also strongly enhanced Zn and copper (Cu) cellular contents (Hoffman and Hardej, 2012). Nevertheless, the same authors also reported that DTC namab, which contains sodium but not Mn, affected Cu and Zn contents in a much lower extent than maneb and mancozeb supporting the involvement of Mn in oxidative stress induced by maneb and mancozeb (Hoffman and Hardej, 2012). In another recent study, authors reported that rats exposed to mancozeb (at 100 mg/kg b. wt. via oral gavage once daily for 28 days) showed no alteration of hepatic histology but a slight increase in Mn and glutathione contents and increases in glutathione peroxidase and reductase activities in the liver (Kistinger and Hardej, 2022). However, in the same study, the use of namab (at 95mg/kg b. wt.) recapitulated the effects of mancozeb on the liver suggesting that the organic backbone of DTCs but not metals (Mn or Zn) was mainly responsible for the alteration of hepatic redox status (Kistinger and Hardej, 2022).

In this context of contradictory data, our study further supports the mechanism of toxicity of the Mn-containing DTCs maneb and mancozeb through their metabolization in hepatocytes generating intracellular Mn accumulation and depletion in zinc (Zn), which are the key events in the observed cytotoxicity. The conclusion that alteration of the Mn/Zn homeostasis plays crucial role in hepatocyte death is based on the following observations 1) exposure to ETU did not induce hepatocyte cell death, 2) toxicity of maneb was prevented by a 10-fold excess in ZnCl_2_, which partially restored the Mn/Zn contents indicating that Zn/Mn intracellular flux are predominant in the pesticide-mediated toxicity, 3) the DTC zineb, with chemical structure similar to maneb and mancozeb but containing only Zn, did not induce hepatocyte cytotoxicity, and 4) exposure of hepatocytes to MnCl_2_ produced similar adverse effects than maneb and mancozeb.

While Mn is an essential metal because of its role as co-factor of multiple enzyme-mediated catalytic activities, high Mn overload resulting from occupational overexposure causes the well-described manganism, a neurological syndrome similar to Parkinson’s disease (Kim et al., 2015). Even in non-occupationally exposure (i.e. lower doses of exposure), Mn remains one of the most toxic metals with endocrine disrupting effects especially during pregnancy with impacts on birth outcomes (Tsai et al., 2015). Mn through its ability to cross the blood-brain barrier more efficiently in infants because of an incompletely developed blood–brain barrier and their inability to fully eliminate this metal, accumulates in the hypothalamus to affect the timing of puberty (Dees et al., 2017).

The characterization of Mn homeostasis was recently established with the identification of the membrane transporters DMT1 (*SLC11A2*), ZIP18 (*SLC39A8*), *SLC30A10* and ZIP14 (*SLC39A14*) regulating Mn uptake and efflux in enterocytes, cholangiocytes and hepatocytes (Katz and Rader, 2019). Importantly, these transporters also regulate transport of other metals including Zn, which is consistent with our data showing that excess in ZnCl_2_ abolished the deleterious effects of DTCs and MnCl_2_ in hepatocytes. This observation may be of interest in order to prevent adverse effects of Mn-containing DTCs during occupational exposure through a specific diet containing Zn supplementation.

The principal risks of exposure to DTCs are expected to be inhalation resulting from spray drift from application sites (van Wendel de Jood et al., 2014) and residues present on agriculture products (EFSA, 2017). Despite the fact that most DTCs will soon be banned in E.U., chronic exposure of worldwide populations to maneb and manozeb at low doses is likely to persist in the future and the impact of both ETU and Mn as endocrine disruptors needs to be considered especially for individuals with liver diseases. In healthy individuals, excess of Mn in blood or liver is released into the bile and eliminated with the stool (Katz and Rader, 2019). Patients suffering from liver diseases could be more prone to Mn toxicity since their Mn removal system through the production and flow of bile may be impaired. Consistently, patients with liver cirrhosis have increased blood and brain Mn contents that may contribute to the occurrence of neurodegenerative disorders (Mehkari et al., 2020). In this context and in the light of our data, the impact of long-term exposures to low doses of Mn-containing DTCs especially in the liver remains to be investigated.

## 5. Conclusion

This work provides new insight into the toxicity of manganese Mn-containing DTCs maneb and mancozeb in metabolically competent human hepatocytes. Very low doses of maneb and mancozeb induce oxidative stress triggering caspase-dependent apoptosis. The mechanism of cytotoxicity of Mn-containing DTCs involves their metabolization and the release of Mn leading to intracellular Mn overload and depletion in zinc (Zn). Alteration of these metal’s homeostasis provokes oxidative stress and apoptosis, which both can be prevented by Zn supplementation.

## Abbreviations

TCs: Dithiocarbamates
CYP450: cytochrome P450
DMSO: dimethyl sulfoxide
XMEs: xenobiotic metabolizing enzymes
HPLC-UV: High Pressure Liquid Chromatography-Ultra Violet
LC-MS: Liquid-Chromatography-mass spectrometry
ROS: Reactive Oxygen Species

## Acknowledgements

This work was funded by the Cancéropole Grand-Ouest (projet structurant Pesticides and Tumor Niches-PeNiCa, 2016-2018, coordinator O. Herault, Tours, France), the Institut National de la Santé et de la Recherche Médicale (Inserm, France) and the project PESTIFAT (Agence Française pour la Biodiversité (AFB), plan ECOPHYTO II driven by the Ministères en charge de l’Environnement et de l‘Agriculture, Appel à projet de Recherche en Environnement-Santé-Trvail (PREST 2018, coordinator B. Fromenty, Rennes, France). Kilian Petitjean received a PhD fellowship from the Agence Française pour la Biodiversité (AFB). The authors would like to thank the Centre de Ressources Biologiques (CRB) Santé of Rennes for providing human hepatocytes as well as Professor Pascal Reyner and Naïg Guegen (Pôle de Recherche et d’Enseignement en Médecine Mitochondriale-PREMMi, Centre Hospitalier Universitaire, Angers, France). Information about protocols and Supporting Information repositories will be publicly available at Mendeley Data website if the manuscript is accepted.

## Author Contributions

Conceptualization, P.LO., A.C. and B.F.; methodology, K.P., S.B., C.R., M.R., O.L., L.A., N.B. and P.LO.; validation, L.A., N.B. and P.L.; formal analysis, P.LO., B.F. and A.C.; investigation, K.P., Y.V., S.B., C.R., C.A., P.LE.; resources, L.A., N.B. and M.R.; data curation, A.C. and P.LO.; writing—original draft preparation, P.LO.; writing— review and editing, K.P., A.C., B.F., N.B.; supervision, P.LO., A.C. and B.F.; project administration, P.LO, A.C., B.F.; funding acquisition, P.LO., C.O., O.H., A.C. and B.F. All authors have read and agreed to the published version of the manuscript.

## Data Sharing

Information about data sharing protocols, options for accessing data, and links to data repositories may be provided in the “Acknowledgments” section, as noted above. Authors may also provide links to data repositories in the “Methods” or “Results” sections of their manuscripts, as appropriate.

