## Supplemental Figures 1 to 5 for "Low concentrations of ethylene bisdithiocarbamate pesticides maneb and mancozeb impair manganese and zinc homeostasis to induce oxidative stress and caspase-dependent apoptosis in human hepatocytes"

#### Supporting Information 1

##### *Pesticides:*

**Table**

| | | %ADI | | | ADI | | |
|---|---|---|---|---|---|---|---|
| **Pesticides** | **CAS Number** | **(µg/kg/days)** | **(µg/L)** | **(µM)** | **(mg/kg/days)** | **(mg/L)** | **(µM)** |
| Chlorpyrifos | 2921-88-2 | 0.292 | 3.51 | 0.01 | 0.01 | 0.12 | 0.3423 |
| Diazinon | 333-41-5 | 0.076 | 9.13 | 0.03 | 0.0002 | 0.0024 | 0.0078 |
| Dimethoate | 60-51-5 | 0.191 | 2.29 | 0.01 | 0.001 | 0.012 | 0.0523 |
| Imazalil | 35554-44-0 | 1.734 | 20.80 | 0.07 | 0.025 | 0.3 | 1.009 |
| Iprodione | 36734-19-7 | 0.275 | 3.30 | 0.01 | 0.06 | 0.72 | 2.181 |
| Maneb | 12427-38-2 | 16.14 | 193.67 | 0.73 | 0.05 | 0.6 | 2.262 |
| Mancozeb | 8018-01-7 | 15.99 | 191.88 | 0.72 | 0.05 | 0.6 | 2.251 |

*References, chemical structures and values of ADI of each pesticide of the mixture and the highest Nutritional Daily Intake (hNDI) or Theorical Maximum Daily Intake (TMDI) further referred as %ADI for French population as given by the 2014 annual report EFSA. All %ADI values for the seven compounds are < to ADI values.*

##### *Chemical structures:*

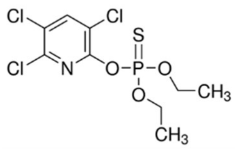

**Chlorpyrifos**

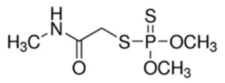

**Dimethoate**

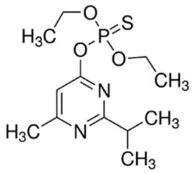

**Diazinon**

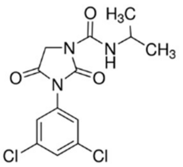

**Iprodione**

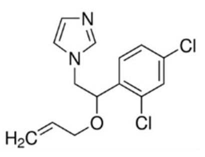

**Imazalil**

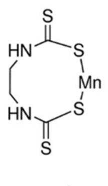

**Maneb**

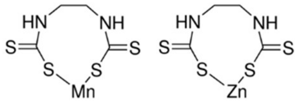

**Mancozeb**

##### *Cell cultures:*

Primary normal BM-MSCs were isolated from healthy donors (without any hematological disorder) undergoing orthopedic surgery (University Hospital, Tours, France) after informed consent and following a procedure approved by the local ethical committee. BM-MSCs were amplified as previously described (Delorme B Met Mol Biol 2007). MSCs (used between passages 4 and 7) were cultured at 2,000 cells per cm2 in α-MEM modified with ribonucleosides and deoxyribonucleosides and supplemented with FCS (10%) and 2 mM L-glutamine, 100 U/ml penicillin, 100 µg/ml streptomycin, and 0.2 ng/ml of FGF2.

Peripheral blood mononuclear cells were obtained from two human buffy coat (Etablissement Français du Sang, Rennes, France) by differential centrifugation on UNI-SEP maxi U10 (Novamed, Jerusalem, Israel). The experiments were performed in compliance with the French legislation on blood donation and blood products' use and safety. Monocytes from healthy donors were enriched using a human CD14 separation kit (Microbeads; Miltenyi Biotec, Bergisch Gladbach, Germany), plated at a density of 0.5 x $10^6$ cells per well in 24-well plates and cultured at 37 C with 5% humidified $CO_2$ in RPMI 1640 medium supplemented with 100 IU/ml penicillin–100 mg/ml streptomycin, 2 mM L-glutamine and 10% FCS BioWhittaker®. Macrophages were obtained after differentiation from monocytes by incubation with 50 ng/mL rhGM-CSF in RPMI 1640 medium during 7 days, as previously described (Vène et al., *Int J Pharm* 2016). All cells were cultured at 37°C in a 5% $CO_2$ atmosphere.

HepG2 and Huh7 human hepatoma cell line were cultured in Dulbecco's Modified Eagle Medium (DMEM) supplemented with 10% FBS, 1% glutamine (200mM), 100U/ml penicillin, 100µg/ml streptomycin.

Rat liver epithelial cells (RLEC) were obtained as previously described from rat livers (Corlu et al., *J Cell Biol* 1991) and maintained in William's E medium supplemented with 10% Fetal calf serum, 100 units/mL penicillin, 100µg/mL streptomycin, 2mM L-glutamine.

***List of forward and reverse primers used for RT-qPCR:***

Supporting Information 2

| | Primer sequence | |
|---|---|---|
| | Forward | Reverse |
| BAX | 5'-GGA-GCT-GCA-GAG-GAT-GAT-TG-3' | 5'-AGT-TGA-AGT-TGC-CGT-CAG-AA-3' |
| DDIT3 | 5'-GTC-TAA-GGC-ACT-GAG-CGT-ATC-3' | 5'-CAC-TTC-CTT-CTT-GAA-CAC-TCT-CT-3' |
| GPX3 | 5'-CAA-CCC-TTT-CTC-TCC-AGT-TCT-C-3' | 5'-CCC-AAG-GTT-GAG-GTA-TCA-GTT-AG-3' |
| SOD1 | 5'-GAA-AAC-ACG-GTG-GGC-CAA-AG-3' | 5'-GCA-GTC-ACA-TTG-CCC-AAG-TC-3' |
| SOD2 | 5'-TTG-CTG-GAA-GCC-ATC-AAA-CG-3' | 5'-GAA-ACC-AAG-CCA-ACC-CCA-AC-3' |
| TBP | 5'-GAG-CTG-TGA-TGT-GAA-GTT-TCC-3' | 5'-TCT-GGG-TTT-GAT-CAT-TCT-GTA-3' |

***Genotoxicity assay using γ-H2AX staining:***

The γ-H2AX staining was used to monitor genotoxicity. After 24 hours of treatment, cells were washed with PBS and then fixed with 4% PAF for 15 minutes at RT. Cells were permeabilized with PBS containing 0.1% Triton X-100 for 5 minutes at RT, rinsed twice and then blocked with FBS 10% in 0.1% PBS-Tween 20 for 1h at RT. Cells were washed twice with PBS 0.1% Tween 20 and then incubated overnight at 4°C with Anti-gamma H2A.X (phospho S139) antibody [EP854(2)Y] (Alexa Fluor 488) ab195188 (abcam) at 1/50 dilution or with non-immune control antibobies labeled with Dylight 488 at 1/250 dilution. Antibodies were diluted in PBS 0.1% Tween 1% BSA. After two washes, nuclear DNA staining was performed using Hoechst 33342 at 10µg/mL in PBS 0.1% Tween 20. Then, cells were washed with PBS and γ-H2AX positive foci were observed by fluorescence microscopy using Zeiss Inverted Microscope and photographs were analyzed with the AxioVision Software.

***Detection of cells with sub-G1 DNA content by flow cytometry:***

Apoptotic cells with hypoploid DNA content were detected by flow cytometry using propidium iodide. Briefly, after 24 hours of treatment, cells were collected and DNA was labeled using the BD Cycletest Plus DNA Kit (BD Bioscience ref: 340242). The fluorescence intensity of 10,000 cells was analyzed with a Becton Dickinson le LSRFortessa™ X-20 (cytometry core facility of the Biology and Health Federative research structure Biosit, Rennes, France).

**Buffers and substrates used for measurement of mitochondrial OXPHOS complex enzyme activities:**

**Table**

| Complex | I | II | III | IV | V | CS |
|---|---|---|---|---|---|---|
| Potassium dihydrogen phosphate | 80mM pH 7.4 | 50mM pH 7.5 | 10mM pH 7.8 | - | 20mM pH 8.1 | - |
| Adjuvants | BSA 1mg/mL | BSA 1mg/mL<br>EDTA 0.1mM | BSA 1mg/mL<br>EDTA 2mM<br>Lauryl maltoside 0.6mM | BSA 1mg/mL<br>Lauryl maltoside 2.5mM | $MgCl_2$ 1mM<br>KCl 2mM<br>Pyruvate kinase 1U/mL<br>LDH 1.7U/mL<br>PEP 0.4mM<br>FCCP 0.15mM | Oxaloacetate 0.5mM<br>Triton X100 0.1% |
| Inhibitors | KCN 1mM<br>Azide 2mM<br>Rotenone 10 µM | KCN 2mM<br>Rotenone 10µM<br>Antimycin 2µM | KCN 0.24mM<br>±Antimycin 4µg/mL | | Antimycin 0.1µg/mL<br>±oligomycine 0.01µg/mL | |
| Susbtrats | Decylubiquinone 0.1m<br>DCPIP 75µM<br>NADH 30µM | Succinate 20mM<br>DCPIP 80µM<br>Decylubiquinone 60µM | Cytochrome C 40µM<br>Decylubiquinol 0.1mM | reduced Cytochrome C 50µM | NADH 0.15mM<br>ATP 0.2mM | DTNB 0.15mM<br>ACoA 0.3mM |

BSA : bovine serum albumin ; DCPIP : 2,6-dichlorophenolindophenol ; KCl : potassium chloride ;; KCN : potassium cyanide ; LDH : lactate dehydrogenase ; $MgCl_2$ : magnesium chloride ; PEP : phosphoenolpyruvate

#### Supporting Information 2

**Metabolism of mancozeb in hepatocyte-like HepaRG cells.**

Based on bibliographic data (references below), 12 metabolites that can be formed during the catabolism of mancozeb have been inventoried. Among these 12 compounds, eight have analytical standards that were purchased and analyzed in this study (Table 1). Ethylene thiourea (ETU) (99,6% purity), mancozeb (70.1%), EDA (100%), N-acetyl-EDA (100%), allantoin (100%), glycine (100%) and creatinine (99.1%) were purchased from Dr. Ehrenstorfer Gmbh, EBIS (100%) from Toronto Research Chemicals. Formic acid, methanol and water were HPLC/MS-grade, and were purchased from J.T. Baker (Paris, France).

The analysis was performed by direct injection in UPLC/MSMS with a Xevo-TQD ® triple quadrupole (Waters) in MRM (Multiple Reaction Monitoring) mode. The mode of ionization retained is positive electrospray.

Chromatographic separation was done with a Waters Acquity UPLC HSS-T3 C18 column (2.1mm x 150 mm, particle size 1.8 µm, Waters®) maintained at 30 °C. The mobile phase was a gradient of water/0.01% formic acid (v/v) (A) and methanol /0.05% formic acid v/V (B) with a flow rate of 0.2 mL/min. The elution gradient started with 95% eluent A, maintaining isocratic conditions for 2 minutes. Then eluent B increased to 100% in 7 minutes and was maintained for 2.50 minutes. Finally, initial conditions were reached again in 0.10 minutes with a re-equilibrium time of 3.4 minutes to restore the column. The injection volume was 5 µL.

For the mass detector, the parameters were as following: Source temperature 150 °C, desolvatation temperature 650 °C, cone gaz flow 50 L/h, desolvation gas flow 800 L/h, spray voltage 1kV.-

For each compound, distinctive ions and transitions were selected for identification (MRM2) and quantification (MRM1), as well as for confirmation purposes.

Dimethyl-sulfoxide (DMSO) was the solvent for ETU, mancozeb, hypoxanthine to prepare stock solution at 1mM and a mixture of water and 0.1% NAH4OH for xanthine and uric acid.

The injection of analytical standards (at a concentration of 10 µM in water) allowed to define the analytical features of the 8 metabolites and active substance (table 1) and to estimate their limits of quantification. 9 concentrations (0.01 to 10 µM in a mixture water/ William's E Medium 50/50 v/v)) were injected to determine the calibration function. Data was best fitted with a quadratic calibration except for ETU (cubic function). A control (5 µM) was injected every 15 samples.

This method was applied to samples, in culture media containing or not mancozeb and incubated or not with HepaRG cells with direct injection (5µL).

| Mancozeb / metabolites | CAS n° | Rt (min) | Cv (V) | Precursor ion | MRM1 | MRM2 | LoQ (µM) |
|---|---|---|---|---|---|---|---|
| Mancozeb | 8018-01-7 | 8.7 | 40 | 213 | 213>86 (25) | 213>169 (17) | 0.01 |
| Ethylenediamine (EDA) | 107-15-3 | 1.5 | 20 | 61 | 61>44 (30) | - | 11 |
| Glycine | 56-40-6 | 1.6 | 22 | 76 | 76>30 (8) | 76>48 (8) | n.d. |
| N-acetyl-EDA (N-(2-aminoethyl)acetamide) | 1001-53-2 | 1.6 | 20 | 103 | 103>44 | 103>43 | 11 |
| Allantoin | 95-59-6 | 1.8 | 25 | 157 | 157>97 (12) | 157>114 (10) | 3 |
| Creatine | 57-00-1 | 1.8 | 30 | 132 | 132>44 (16) | 132>90 (12) | 0.1 |
| EU – ethylene urea | 120-93-4 | 2.6 | 45 | 87 | 87>44 (12) | 87>42 (15) | 0.2 |
| ETU – ethylene thiourea | 96-45-7 | 3.0 | 30 | 103 | 103>44 (14) | 103>60 (21) | 0.1 |
| EBIS (ethylene bisisothiocyanate sulfide) | 33813-20-6 | 8.9 | 40 | 177 | 177>43 (22) | 177>101 (20) | 0.1 |

Table 1 : Detection parameters of mancozeb and its metabolites (Rt: retention time; Cv: cone voltage; MRM1 quantification (uma) (collision energy - eV); MRM2 confirmation (uma) (collision energy - eV); LoQ:Limit of Quantification; n.d.: not determined)

Analyses showed that EU, EDA, N-Acetyl-EDA, allantoin and EBIS were absent of all samples analysed. Glycine was found in all media at nearly identical amounts because this amino acid is a component of the Williams E medium used for the culture of HepaRG cells (concentration in the medium 0.66 mM). Creatine is also found in culture media incubated with cells regardless of the incubation with mancozeb indicating that creatine is most likely produced by cells. Ethylene thiourea (ETU) was found only in media containing mancozeb, and surprisingly, even in absence of cells. The amounts of ETU did not change in culture media of cells that do not metabolize the mancozeb demonstrating that the ETU found in these media was a degradation product present in the stock solution of mancozeb. In contrast, the amounts of ETU increased in the culture media of hepatocyte-like HepaRG cells exposed to mancozeb demonstrating the catabolism by these cells of this fungicide into its main metabolite.

For the 5 other metabolites of mancozeb (2-imidazoline, ethylene thiuram disulfide (ETD), ethylene thiuram monosulfide (ETM), ethylenethiourea-N-thiocarbamide (ETT) and Jaffe's base sulfoxide) for which it was not possible to obtain analytical standards, a full scan screening was realised and their mass were scrutinized. No peak with these exact masses were found in the samples, which confirmed their absence in the culture media of HepaRG cells incubated with mancozeb.

**Identification of xanthine.**

Incubation of mancozeb with hepatocyte-like HepaRG cells led to the production of two compounds detected by HPLC/UV at 247 nm at retention times of 2.7 and 4.5 minutes. The peak detected at 2.7 minutes co-eluted with ETU standard (data not shown) and was confirmed as ETU by UPLC-MS/MS.

To identify this compound, the first step was to determine its exact mass. To do this, the samples were injected onto an UPLC/MSMS with a Xevo-TQD ® triple quadrupole (Waters) in MRM (Multiple Reaction Monitoring) mode under analytical conditions similar to those used in HPLC in order to maintain a similar elution order of compounds and retention time.

The masses observed in positive electrospray ionization mode (153) and in negative ionization mode (151) led to the conclusion that the compound has a molecular mass of 152 g/mol. The use of the MASS BANK database (public mass spectrum library referencing about 10,000 compounds: https://massbank.eu/MassBank/Search) made it possible to identify two potential candidates: xanthine (CAS 69-89 -6) and oxypurinol (CAS 2465-59-0), two compounds of the purine family. We purchased the analytical standards (Tokyo Chemical Industry Co) of these two compounds (Oxypurinol - 95.1% / xanthine – 100 %) and their injection allowed the identification of the unknown compound: xanthine (Table 2).

The presence of xanthine was confirmed in the culture media of control cells unexposed to mancozeb at levels approximately 10 times lower than the amounts detected in media of cells incubated with 100 □M mancozeb. Importantly, we found that exposure of HepaRG cells to 100 □M $MnCl_2$ led to the production and release of xanthine in culture media in the same amounts as those found in cultures exposed to mancozeb demonstrating that xanthine was not a metabolite of mancozeb catabolism but a compound produced in response to both mancozeb and $MnCl_2$ treatment (**Figure 5**, **Table 1**).

| Putative compounds based on molecular mass | CAS n° | Rt (min) | Cv (V) | Precursor ion | MRM1 | MRM2 | LOQ (µM) |
|---|---|---|---|---|---|---|---|
| Xanthine | 69-89-6 | 4.5 | 40 | 153 | 153>136 (24) | 153>110 (16) | 0.10 |
| Oxypurinol | 2465-59-0 | 5.0 | 40 | 153 | 153>80 (26) | 153>136 (24) | 0.50 |

Table 2 : Parameters for the identification of Xanthine and Oxypurinol in positive electrospray ionization mode (Rt: retention time; Cv: cone voltage; MRM1 quantification (uma) (collision energy - eV); MRM2 confirmation (uma) (collision energy - eV); LoQ: Limit of Quantification)

Considering the biosynthesis and catabolism of the xanthine, we decided to also analyze hypoxanthine and uric acid, two main compounds of this metabolic pathway. Analytical standard hypoxanthine (99.5%) were purchased from HPC standards and uric acid (100%) from Sigma-Aldrich and were injected to determine their retention time and their MS/MS parameters for identification and quantification. We also optimized the analytical method previously developed to quantify mancozeb, ETU, xanthine, hypoxanthine and uric acid (Table 3), by adjusting the elution parameters and cone voltage for the mass detector. An example of a chromatogram is shown in Figure 1. A mixture of water and 0.1% NAH4OH was the solvent for prepare a stock solution of xanthine and Dimethyl-sulfoxide (DMSO) for hypoxanthine.

We found that both xanthine and hypoxanthine were produced by hepatocyte-like HepaRG cells incubated with mancozeb but not uric acid. In addition, we evidenced that both xanthine and hypoxanthine were also produced upon exposure to $MnCl_2$ demonstrating that these compounds are not metabolites of mancozeb.

| Compound | CAS n° | Rt (min) | CV(V) | Precursor ion | MRM1 | MRM2 | LOQ (µM) |
|---|---|---|---|---|---|---|---|
| ETU | 96-45-7 | 2.1 | 30 | 103 | 103>44 (14) | 103>60 (21) | 0.1 |
| Uric acid | 69-93-2 | 2.5 | 40 | 169 | 169>70 (21) | 169>141 (15) | 0.1 |
| Hypoxanthine | 68-94-0 | 2.9 | 25 | 137 | 137>110 (19) | 137>55 (25) | 0.2 |
| Xanthine | 69-89-6 | 3.4 | 35 | 153 | 153>136 (24) | 153>110 (16) | 0.02 |
| Mancozeb | 8018-01-7 | 7.0 | 40 | 213 | 213>86 (25) | 213>169 (17) | 0.05 |

Table 3: Parameters for the identification of compounds, once xanthine has been identified, in positive electrospray ionization mode Rt: retention time; Cv: cone voltage; MRM1 quantification (uma) (collision energy - eV); MRM2 confirmation (uma) (collision energy - eV); LoQ: Limit of Quantification)

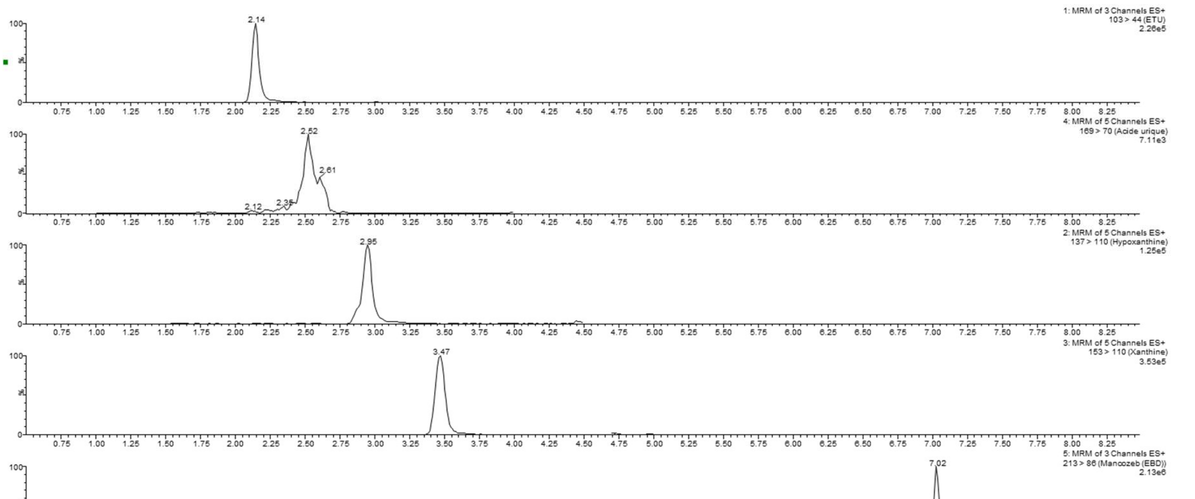


Figure 1 : Chromatogram obtained for an experimental sample (mancozeb 7 µM 3h) (ETU 1.2 µM ; uric acid 0.6 µM ; hypoxanthine 1.35 µM ; xanthine 1.2 ; mancozeb 57µM).

**Chemical structures of some DTCs.**

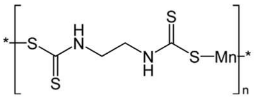

**Maneb**

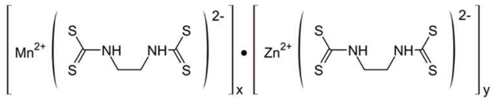

**Mancozeb**

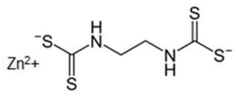

**Zineb**

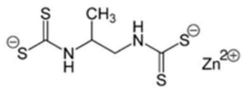

**Propineb**

DTCs share a chemical backbone ($R_2NCS_2R$). Some of them such as maneb, mancozed, zineb and propineb are salts containing $Mn^{2+}$ and/or $Zn^{2+}$. (Ajiboye et al., 2022).

**References**:

World Health Organization. Dithiocarbamate Pesticides, Ethylenethiourea and Propylenethiourea: A general introduction. International programme on chemical safety. Environmental Health Criteria 78. Published under the joint sponsorship of the United Nations Environment Programme, the International Labour Organisation and the World Health Organization. Geneva, 1988

AJIBOYE T.O., AJIBOYE TT, MAEZOUKI R., ONWUDIWE D.C. The Versatility in the Applications of Dithiocarbamates. Int J Mol Sci. 2022, 23(3):1317.

ALDRIDGE, W.N. & MAGOS, L. (1978) *Carbamates, thiocarbamates,* Dithiocarbamates pesticides, ethylenethiourea, and propylenethiourea: a general introduction. Luxembourg, Commission of the European Communities (Report No. V/F/1/78/75 EN).

ENGST, R. & SCHNAAK, W. (1967) [Studies on the metabolism of the ethylenebisdithiocarbamate fungicides maneb and zineb.] *Z. Lebensm. Unters. Forsch.,* **134**(4): 216-221.

ENGST, R. & SCHNAAK, W. (1970a) [Studies on the metabolism of the ethylenebisdiothiocarbamate fungicides maneb and zineb.] *Z. Lebensm. Unters. Forsch.,* **143**: 99-103

FREUDENTHAL, R.I., KERCHNER, G.A., PERSING, R.L., BAUMEL, J., & BARON, R.L. (1977) Subacute toxicity of ethylene bisisocyanate sulfide in the laboratory rat. *J. Toxicol. environ. Health,* **2**:1067-1077.

### Supporting Information 3.

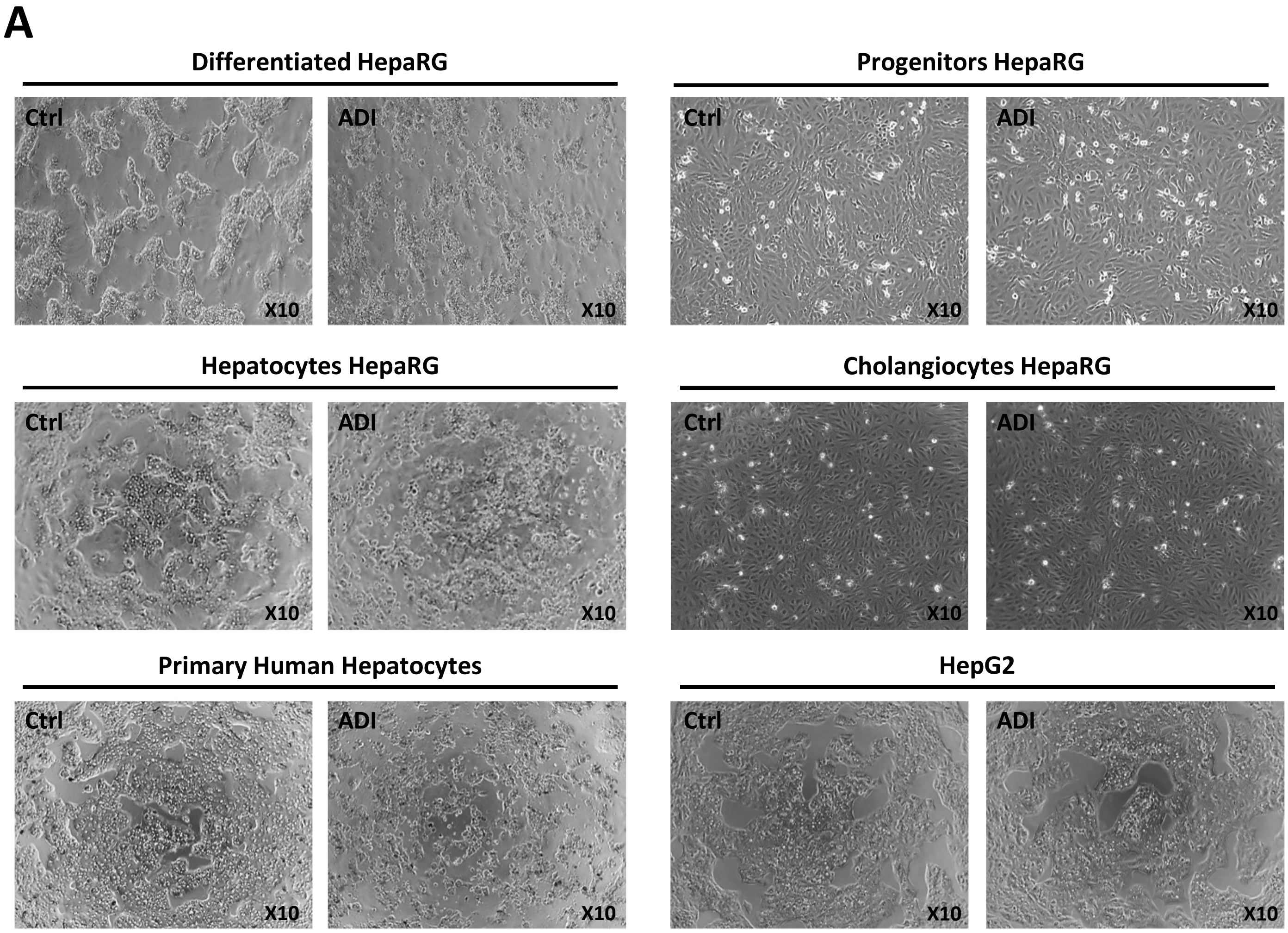

B

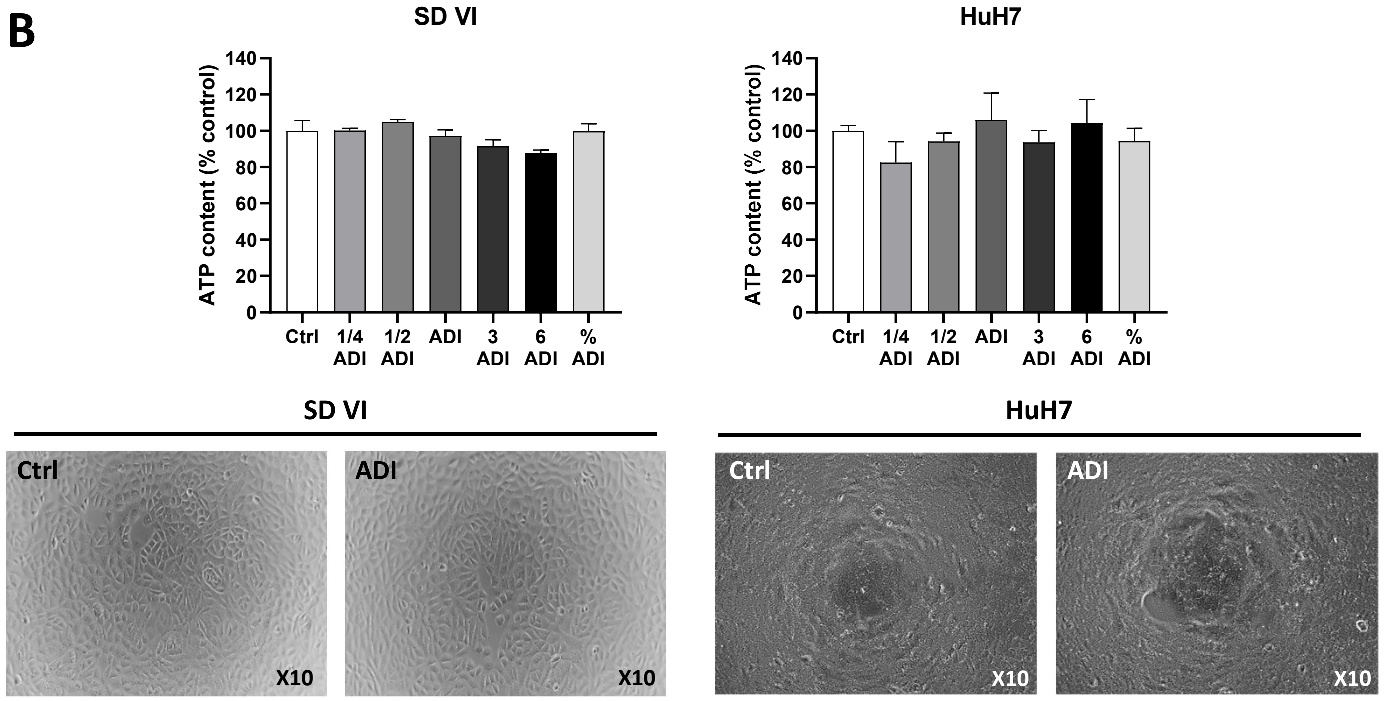
SD VI
HuH7
ATP content (% control)
0
20
40
60
80
100
120
140
Ctrl
1/4 ADI
1/2 ADI
ADI
3 ADI
6 ADI
% ADI
SD VI
HuH7
Ctrl
ADI
X10

C

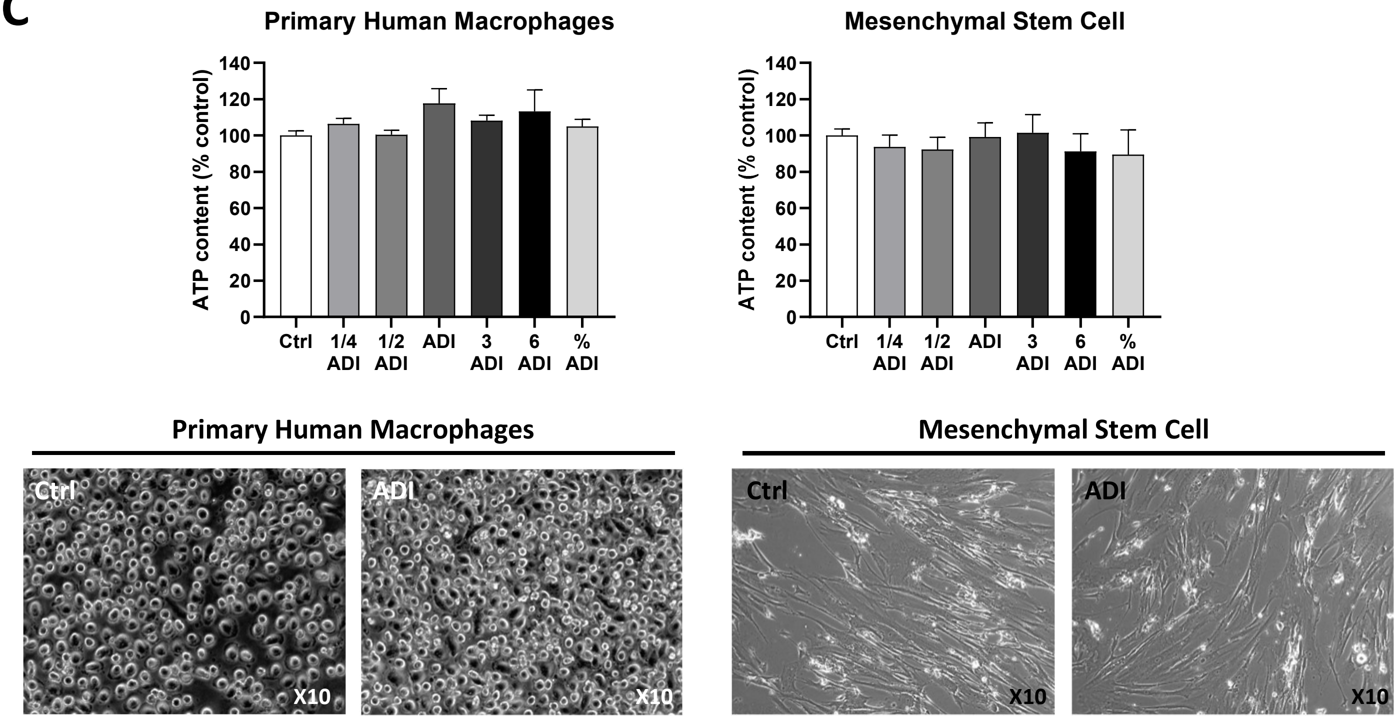
Primary Human Macrophages
Mesenchymal Stem Cell
ATP content (% control)
0
20
40
60
80
100
120
140
Ctrl
1/4 ADI
1/2 ADI
ADI
3 ADI
6 ADI
% ADI
Primary Human Macrophages
Mesenchymal Stem Cell
Ctrl
ADI
X10

**D**

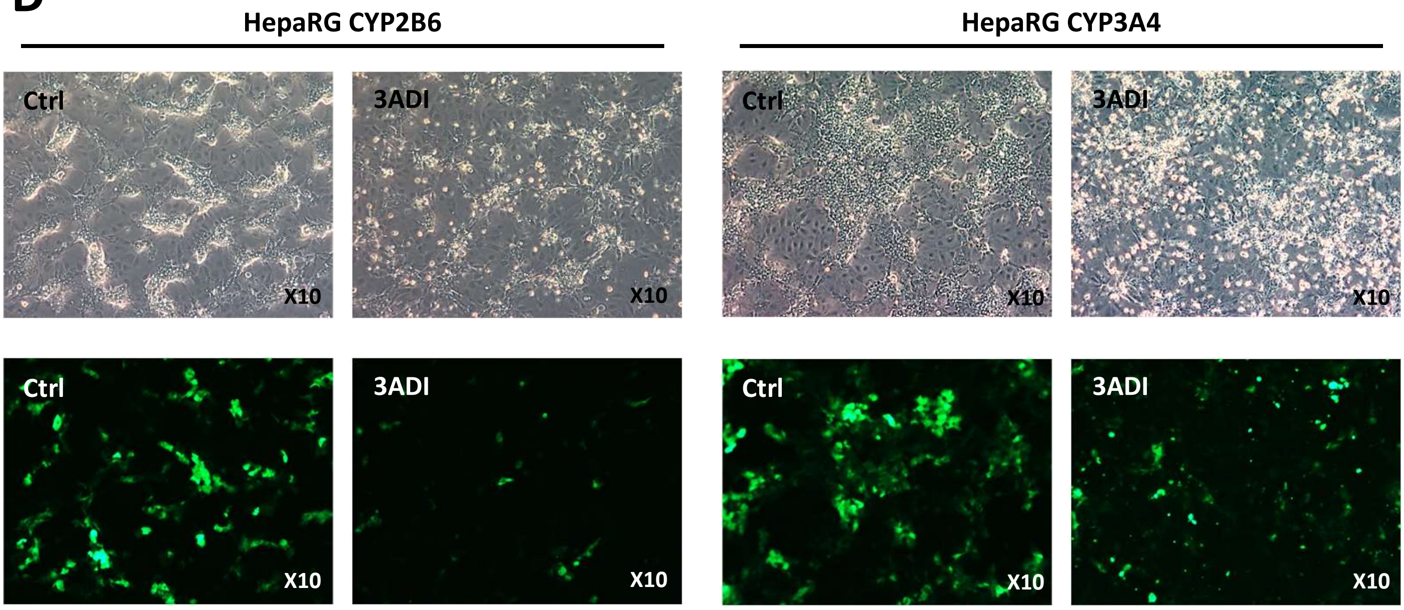


**E**

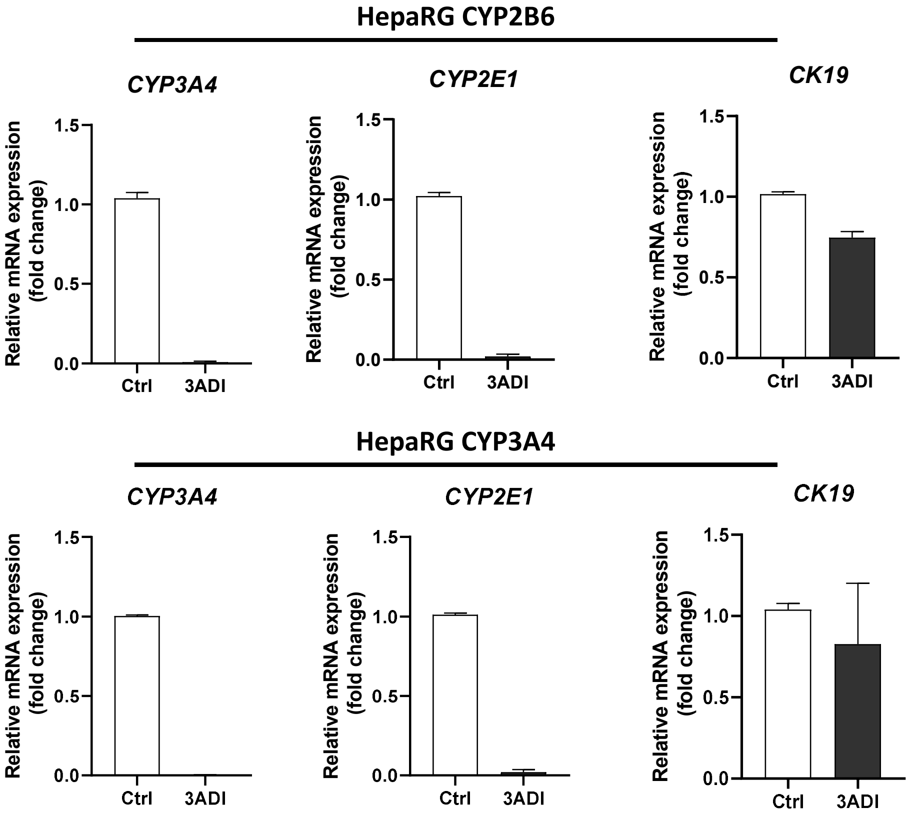


**Supporting Information 3**. **Acute exposure of cell models to the pesticide mixture**.

**A.** Cell morphology in phase contrast microcopy of progenitor and differentiated HepaRG cells (hepatocytes and cholangiocytes), human hepatocytes and HepG2 hepatoma cells in untreated condition (Ctrl) or after 48 hours of exposure with the pesticide mixture (ADI). Cell viability was quantified by measuring the ATP content and cell morphologies in phase contrats microscopy in rat liver epithelial cells (RLEC) SDVI and HepG2 human hepatoma cells (**B**), human macrophages and mesenchymal stem cells (**C**) in untreated control cultures and after 48 hours of exposure to the pesticide mixture at different concentrations (1/4ADI, 1/2ADI, ADI, 3ADI, 6ADI or %ADI). Datas were expressed as % of control. Results are means ± SEM of at least 2 independent experiments. **D.** Cell morphology in phase contrats and GFP detection by fluorescence microscopy in HepaRG-CYP2B6 and HepaRG-CYP3A4 transgenic cells in untreated cultures (Ctrl) and after 48 hours of exposure to the pesticide mixture (3ADI). **E.** Relative expression of *CYP3A4*, *CYP2E1* and *cytokeratin 19* (*CK19*) mRNA levels in untreated control (Ctrl) and pesticide-treated HepaRG-CYP2B6 and HepaRG-CYP3A4 cells.

### Supporting Information 4.

## A

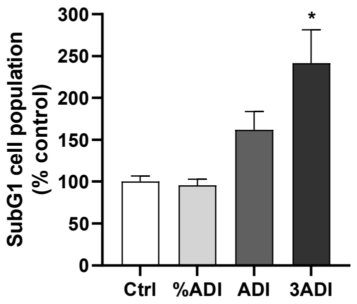


## B

| | I/CS | II/CS | III/CS | IV/CS | V/CS |
|---|---|---|---|---|---|
| **Ctrl** | 0.33 ± 0.02 | 0.45 ± 0.05 | 0.44 ± 0.06 | 1.66 ± 0.35 | 0.95 ± 0.24 |
| **Maneb 6ADI** | 0.35 ± 0.07 | 0.45 ± 0.05 | 0.42 ± 0.12 | 1.70 ± 0.24 | 1.04 ± 0.09 |
| **$MnCl_2$14µM** | 0.30 ± 0.08 | 0.41 ± 0.10 | 0.45 ± 0.07 | 1.73 ± 0.11 | 0.85 ± 0.14 |

## C

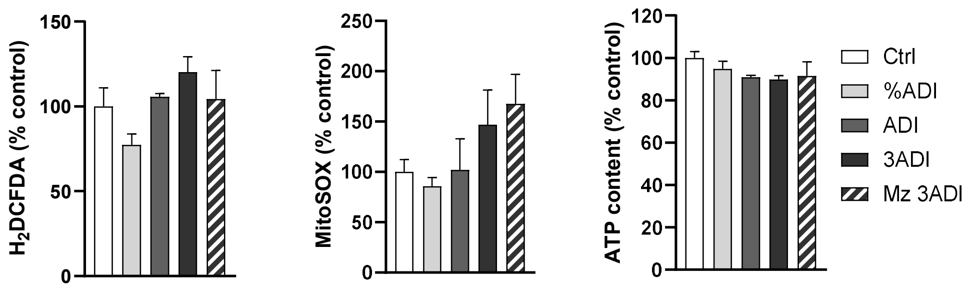


## D

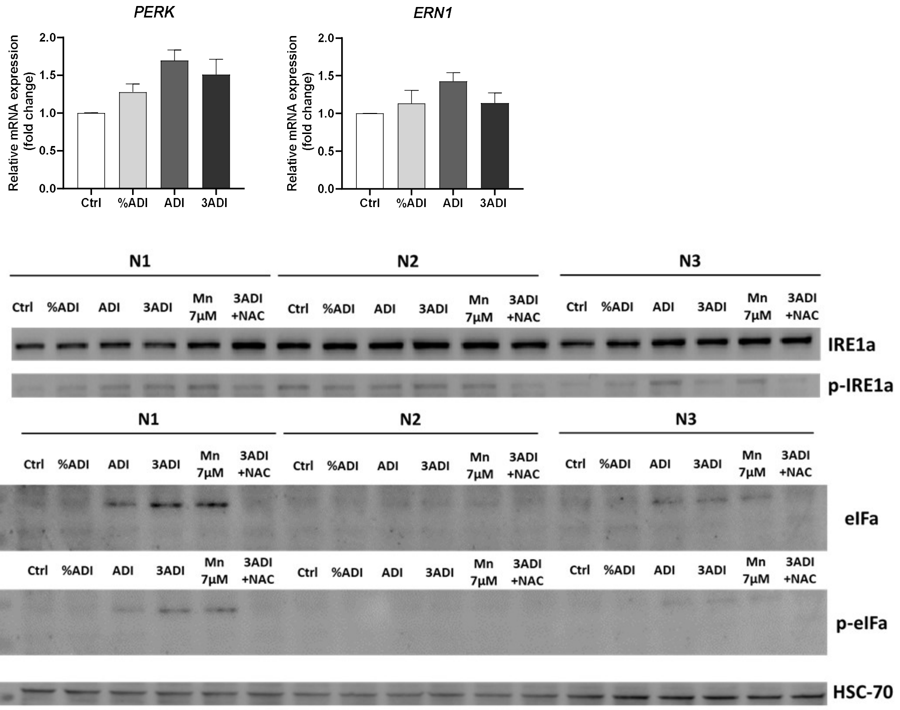

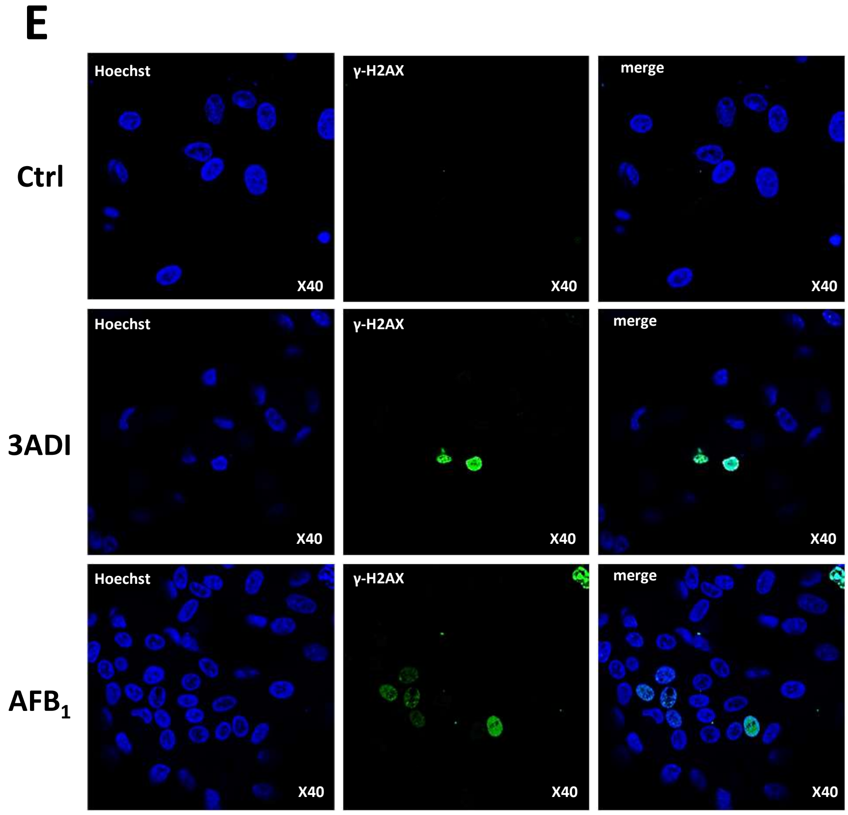


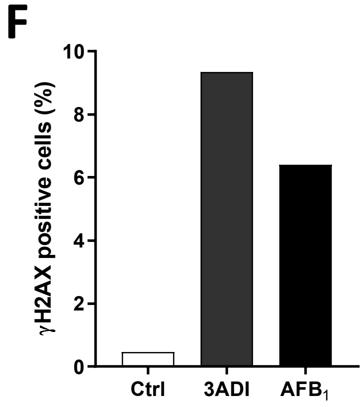


**Supporting Information 4. Apoptosis and genotoxicity in HepaRG cells and human hepatocytes exposed to the pesticide mixture. A.** Percentages of apoptotic HepaRG cells estimated through the quantification of SubG1 population by flow cytometry after exposure to the pesticide mixture for 24h. DNA was stained using propidium iodide. **B**. Activity of the mitochondrial complexes I to V in control culture and in cells exposed to maneb and no significant differences in enzymatic activities could be observed between untreated and pesticide-exposed cells. Enzymes activities of all complex were expressed as nmol substrate/min/$10^6$cells using the Beer Lambert's law and were normalized to the citrate synthase activity (N=3). **C**. ROS production was evaluated using $H_2$DCFDA and MitoSOX probes and cell viability was evaluated by quantification of the relative ATP contents in cholangiocyte-like HepaRG cells exposed for 24h to the pesticide mixture (%ADI, ADI or 3ADI) or mancozeb alone (Mz 3ADI). Data were normalized to the untreated control culture (Ctrl). Results are means ± SEM (N≥3), *$p < 0.05$ significantly different from control (Ctrl). **D**. Relative mRNA levels of *PERK* and *ERN1* (upper charts) and western blot analysis of IRE1$\alpha$ phosphorylation and eIF1$\alpha$ protein expression in hepatocyte-like HepaRG cells (N=3). **E**. Detection of γ-H2AX positive HepaRG cells by fluorescence microscopy in untreated cells (Ctrl) and exposure to the pesticide mixture for 16h (3ADI). Aflatoxin $B_1$ (24h, 0,25µM) was used as genotoxic positive control (AFB$_1$). **F**. Quantification of γ-H2AX positive cells versus total nuclei (Hoechst staining) using Image J. Results are means of at least 3 independent experiments (N≥3).

**Table** Quantities (ng/well) ± SEM of manganese and zinc by ICP-MS

| | | Intracellular | | | Mitochondria | | |
|---|---|---|---|---|---|---|---|
| | | **Hepatocytes** | **Cholangiocytes** | **Progenitors** | **Hepatocytes** | **Cholangiocytes** | **Progenitors** |
| **Manganese (ng/well)** | Ctrl | 12.14 ± 0.43 | 1.72 ± 0.41 | 1.23 ± 0.10 | 0.73 ± 0.10 | 0.46 ± 0.04 | 0.26 ± 0.00 |
| | Ctrl + $ZnCl_2$ 70µM | 14.13 ± 4.34 | | | 0.85 ± 0.04 | | |
| | $MnCl_2$ 7µM | 65.20 ± 15.37 | 6.19 ± 1.98 | 3.06 ± 0.39 | 7.43 ± 2.39 | 0.77 ± 0.08 | 0.37 ± 0.02 |
| | $MnCl_2$ 7µM + $ZnCl_2$ 70µM | 39.78 ± 1.99 | | | 3.34 ± 0.78 | | |
| | Maneb 3ADI | 55.47 ± 8.28 | 4.85 ± 0.69 | 3.51 ± 0.28 | 11.03 ± 7.06 | 0.81 ± 0.19 | 0.46 ± 0.07 |
| | Maneb 3ADI + $ZnCl_2$ 70µM | 39.28 ± 1.85 | | | 3.94 ± 1.33 | | |
| **Zinc (ng/well)** | Ctrl | 122.05 ± 12.78 | 26.76 ± 4.81 | 30.20 ± 1.44 | 13.02 ± 3.27 | 5.04 ± 1.83 | 2.86 ± 0.87 |
| | Ctrl + $ZnCl_2$ 70µM | 140.21 ± 3.08 | | | 4.18 ± 0.79 | | |
| | $MnCl_2$ 7µM | 74.55 ± 11.70 | 43.11 ± 13.96 | 29.44 ± 0.89 | 8.70 ± 2.68 | 9.73 ± 4.08 | 2.03 ± 0.20 |
| | $MnCl_2$ 7µM + $ZnCl_2$ 70µM | 110.96 ± 18.75 | | | 5.17 ± 0.90 | | |
| | Maneb 3ADI | 71.36 ± 6.66 | 23.19 ± 3.51 | 31.48 ± 2.37 | 12.10 ± 4.54 | 4.11 ± 1.06 | 4.33 ± 1.35 |
| | Maneb 3ADI + $ZnCl_2$ 70µM | 111.45 ± 12.09 | | | 6.55 ± 1.72 | | |

**Supporting Information 5**. **Quantification of Mn and Zn in HepaRG cells by ICP-MS.** Resultsexpressed in ng of metals/culture wells are means of at least 3 independent experiments (N≥3).
